# Dystrophin deficiency impairs cell junction formation during embryonic myogenesis

**DOI:** 10.1101/2023.12.05.569919

**Authors:** Elise Mozin, Emmanuelle Massouridès, Virginie Mournetas, Clémence Lièvre, Audrey Bourdon, Dana L Jackson, Jonathan S Packer, Juyoung Seong, Cole Trapnell, Caroline Le Guiner, Oumeya Adjali, Christian Pinset, David L Mack, Jean-Baptiste Dupont

## Abstract

Mutations in the *DMD* gene lead to Duchenne muscular dystrophy, a severe X-linked neuromuscular disorder that manifests itself as young boys acquire motor functions. DMD is typically diagnosed at 2 to 4 years of age, but the absence of dystrophin negatively impacts muscle structure and function before overt symptoms appear in patients, which poses a serious challenge in the optimization of standards of care. In this report, we investigated the early consequences of dystrophin deficiency during skeletal muscle development. We used single-cell transcriptome profiling to characterize the myogenic trajectory of human pluripotent stem cells and showed that DMD cells bifurcate to an alternative branch when they reach the somite stage. Here, dystrophin deficiency was linked to marked dysregulations of cell junction protein families involved in the cell state transitions characteristic of embryonic somitogenesis. Altogether, this work demonstrates that *in vitro*, dystrophin deficiency has deleterious effects on cell-cell communication during myogenic development, which should be considered in future therapeutic strategies for DMD.

## Introduction

Mutations in genes involved in skeletal muscle functions trigger a spectrum of diseases which can lead to significant motor and respiratory impairments, sometimes shortly after birth^1–3^. The cellular and molecular mechanisms of these diseases are complex and result in the breakdown of skeletal muscle homeostasis. In Duchenne muscular dystrophy (DMD) patients, the lack of functional dystrophin causes systemic pathological perturbations, which mostly affect skeletal muscles, heart and the central nervous system. In muscle cells, *DMD* mutations lead to sarcolemmal membrane fragility, calcium overload, oxidative stress, mitochondrial impairments, development of an inflammatory microenvironment and a pro-fibroadipogenic process^4^. As muscles develop in the absence of dystrophin, capturing the pathological molecular cascade downstream of the mutation and tracking progression over time are difficult using animal models. Early studies described prenatal signs of muscle wasting in presymptomatic *mdx* mice and GRMD puppies^5,6^. Signs of muscle damage such as variable myotube diameter, hyaline fibers and internal nuclei had also been observed in fetuses at risk for DMD^7,8^. The most well-characterized functions of dystrophin have been described in fully differentiated skeletal muscle cells, which express the longest isoform of the protein (Dp427m) that connects the contractile machinery with the sarcolemma and the extracellular matrix through the dystrophin-associated protein complex (DAPC)^4^. In other organs, the expression of the *DMD* locus gives rise to other protein isoforms with specific N-terminal regions. How dystrophin expression arises during the embryonic development of different cell lineages remains difficult to characterize with available *in vivo* models. In the embryo, skeletal muscles emerge from the paraxial mesoderm, which differentiates into transient metameric structures called somites, and then into the dermomyotome^9^. This involves a succession of well-orchestrated cell division, migration and transitions between epithelial and mesenchymal states, that later give rise to myogenic progenitors with migratory properties^10–12^. In this context, the expression of dystrophin and the consequences of pathological mutations during skeletal muscle development remain to be characterized in a reliable model.

Pluripotent stem cells represent a proxy for human embryonic development, and they allow the characterization of early disease mechanisms in a dish^13–15^. Several groups have published protocols to differentiate human induced pluripotent stem cells (hiPSCs) into paraxial mesoderm progenitors, somite intermediates, dermomyotome and ultimately skeletal muscle progenitors^16–19^. Recently, our group has used hiPSCs from DMD patients to demonstrate that marked transcriptome dysregulations occur prior to skeletal muscle commitment^20^. Characterization of *DMD* gene expression during early mesoderm induction had led to the identification of an embryonic isoform of dystrophin (Dp412e) in mesoderm progenitors and embryoid bodies^21^. This isoform possesses an alternative exon 1 and encodes an N-truncated protein starting in exon 6 of the skeletal muscle dystrophin isoform (Dp427m). However, its functions and molecular partners remain unknown.

Transcriptomic analysis of hiPSC derivatives at single-cell resolution offers the opportunity to shed light on complex biological processes such as embryonic development and to investigate the effects of disease-causing mutations in any given lineage intermediate. In particular, single-cell RNA-sequencing (scRNA-Seq) has helped characterize the diversity of skeletal muscle cells involved in muscle development^22,23^, specification^24,25^, ageing^26–28^, and in various models of DMD^29–32^. Reconstruction of single-cell trajectories thanks to pseudotemporal ordering of cells has been used to describe dynamics and gene expression profiles during differentiation of myoblasts^33,34^. Studying the divergence of this trajectory when hiPSCs harbor a mutated *DMD* gene will provide a deeper understanding of the molecular drivers of pathology immediately downstream of the mutation and the cascade of molecular events after disease initiation.

In this study, we established the single-cell trajectory of healthy and dystrophin-deficient hiPSCs subjected to myogenic differentiation. We showed that hiPSCs derived from a DMD patient diverged from healthy control cells as early as the somite stage and that the differences propagated as differentiation progressed. After inclusion of an isogenic CRISPR-engineered DMD line, we demonstrated that several cell junction gene families were markedly dysregulated as a consequence of dystrophin deficiency in somite progenitors. Further characterization identified epithelial and mesenchymal populations coexisting *in vitro*, reflecting the cell state transitions occurring in the embryo, and that the formation of epithelial islets is altered in the absence of dystrophin. Finally, we confirmed our results in three additional DMD hiPSC lines by immunostaining and analysis of previous bulk RNA-Seq data generated in our group, which strongly suggests that *DMD* mutations have significant consequences during prenatal development. Overall, this study indicates that dystrophin deficiency leads to disruptions of cell-cell communication and coordinated migration during somitogenesis.

## Results

### DMD hiPSCs bifurcate to an alternative branch of the myogenic trajectory at the somite stage

Human iPSCs from a DMD patient harboring an exon 50 deletion in the *DMD* gene were subjected to directed differentiation into the skeletal muscle lineage in parallel with a healthy control^35^, using a transgene-free and serum-free protocol^16^. Microscopic monitoring showed extensive cell proliferation and densification into multilayered cultures over time (Figure S1A). The differentiation was stopped at Day 28 when fields of thin and spindly myotubes could be observed in the healthy control line. To investigate the dynamics of myogenic differentiation and the impact of dystrophin deficiency, cells were isolated at ten discreet time points from Day 0 (D0) to Day 28 (D28), tagged with barcoded primers during *in situ* reverse-transcription and pooled for single-cell combinatorial indexing RNA-Seq (sci-RNA-Seq, Figure 1A)^36^. A total of 1917 individual cells could be retrieved from 20 distinct samples corresponding to the 2 hiPSC lines across 10 time points (Table S1), and used in the subsequent clustering analyses. After Uniform Manifold Approximation and Projection (UMAP) for dimension reduction, cells were distributed among 7 clusters expressing well-defined marker genes associated with pluripotency at Day 0 (*e.g. POU5F1*, clusters 1-2), primitive streak at Day 2 (*T,* cluster 5), paraxial mesoderm at Day 2-4 (*TBX6*, cluster 3), somite and dermomyotome at Day 7-10 (*PAX3*, clusters 2-7), and ultimately skeletal muscle progenitor cells from Day 22-28 (*PAX7*, *MYOD1*, cluster 1) (Figure S1B-D). Interestingly, DMD and healthy control cells were intermingled into superimposed clusters during the first week (Day 0 – 7), but from Day 10, they started to form distinct cell populations (Figure 1B-C). Single-cell trajectory reconstruction with Monocle^34^ identified a bifurcation point between Day 7 and Day 10, from which most of the cells segregated on two distinct branches in a genotype-dependent manner (Figure 1D-E). More precisely, 86 % of the healthy cells after the divergence were on a single branch (621 vs. 103), while 86 % of the DMD cells were on the other branch (329 vs. 55, Figure S1E-F). Branch expression analysis modelling (BEAM) identified thousands of differentially-expressed genes (DEG) along the WT and the DMD-enriched branches. The most significant candidates (1987 genes with p-adj < 0.0001) were clustered based on their expression dynamics on the two branches for gene ontology analysis (Figure 1F, Table S2). Interestingly, modules of genes overexpressed along the healthy branch were enriched for terms related to muscle development (*e.g.* muscle cell differentiation, muscle contraction, actin cytoskeleton organization). In contrast, gene modules overexpressed along the DMD branch matched with GO terms related to the development of alternative lineages, particularly neurons (*e.g.* neurogenesis, synapse organization, axon guidance). To gain insight into the regulation of myogenesis in pseudotime, differential expression of skeletal muscle markers was assessed between the two branches. Significant differences in pseudotemporal dynamics were found for critical regulators of myogenesis, including *MYOD1*, *MYOG*, *MEF2C*. Notably, the master regulators *MYOD1* and *MYOG* were expressed at very low levels in DMD cells, in contrast with *PAX7* and *MEF2C*, although for the latter, expression in DMD cells was significantly reduced as pseudotime progressed (Figure 1G). In addition, genes coding for important structural proteins such as *MYH3*, *MYH8*, *DES* or *TTN* were also found significantly dysregulated (Figure 1G). Thus, hiPSCs derived from a DMD patient deviated from the myogenic trajectory followed by healthy control cells at the somite stage, resulting in dysregulated expression of myogenesis regulators.

**Figure 1:**
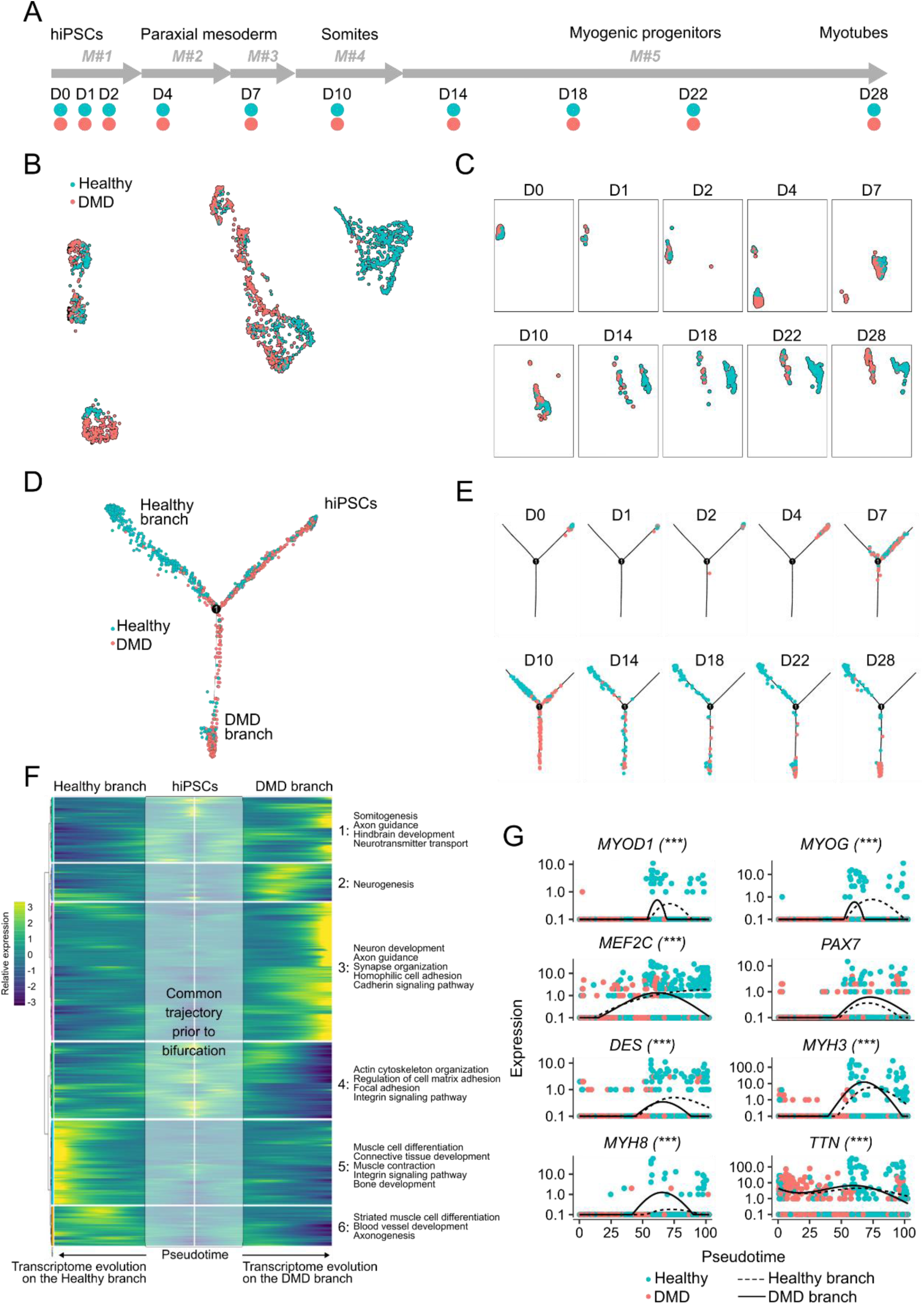
Myogenic differentiation of DMD and healthy control hiPSCs at the single-cell resolution. (A) Cell collection timeline along hiPSC myogenic differentiation using the combination of 5 defined media (M#1 to M#5) described previously^16^. D0 to D28: Day 0 to Day 28. (B) UMAP plot showing the 1917 individual cells colored by hiPSC line of origin. (C) Deconvolution of the UMAP plot by collection time point (D0 to D28). (D) Single-cell trajectory capturing the dynamics of myogenic differentiation as DMD and Healthy hiPSCs progress along the initial common branch (hiPSCs) to the bifurcation point (1) and one of the two alternative branches. (E) Deconvolution of the single-cell trajectory by collection time point (D0 to D28). (F) Branched expression analysis modeling identifying modules of genes with branch-dependent expression. Key Gene Ontology terms associated with each individual module are indicated on the right-hand side. (G) Gene expression kinetic in pseudotime for myogenic markers along the Healthy (dotted line) and the DMD (full line) branch. Each dot shows an individual cell with its computed pseudotime value. Significant differences in branch-dependent expression are indicated with *** (p-adj < 0.001).

### DMD patient-derived hiPSCs exhibit a marked dysregulation of cell junction genes at Day 10

As the deviation of DMD cells was first evident at Day 10 on the single-cell trajectory, “Day10” cells were reanalyzed separately. Although a single UMAP cluster was observed, DMD and healthy control cell were clearly separated (Figure 2A). Differential expression analysis using the regression model from Monocle 3 identified 94 genes significantly dysregulated in DMD cells (adjusted p-value < 0.01) (Table S3). Among these differentially expressed genes (DEGs), multiple cell junction and extracellular matrix genes were identified, including cadherins and proto-cadherins (*CDH11*, *PCDH9*), integrins (*ITGA1*, *ITGA4*, *ITGB1*, *ITGAV*), fibronectin (*FN1*) and numerous collagens (*COL1A2*, *COL3A1*, *COL4A1*, *etc.*). Gene ontology (GO) analysis confirmed significant enrichments in related biological processes, such as cell-matrix interaction (p = 5.6E-4), cell-cell adhesion (p = 6.9E-7), extracellular matrix organization (p = 6.9E-4) and cell junction organization (p = 4.0E-2) (Figure 2B). Strikingly, PANTHER Pathway statistical enrichment test only identified 3 pathways as significantly overrepresented in the DEG list, and two of them were directly related to specific cell junction protein families: the integrin and cadherin signaling pathways (p = 2.4E-12 and p = 4.2E-2, respectively) (Figure 2C). Of note, the GO analysis also indicated that DEGs included regulators of key developmental processes in non-muscle lineages, such as neurogenesis and axon guidance (*SEMA3A*, *EPHA3*, *NRP1*, *NEFL*, *UNC5C*), and angiogenesis (*ANGPT1*, *THSD7A*). Neuronal by-products have already been observed when hiPSCs are differentiated with this specific protocol^37^. In this study, the DMD hiPSC line might be more prone to such uncontrolled differentiation events, leading to the formation of more alternative cell types.

**Figure 2:**
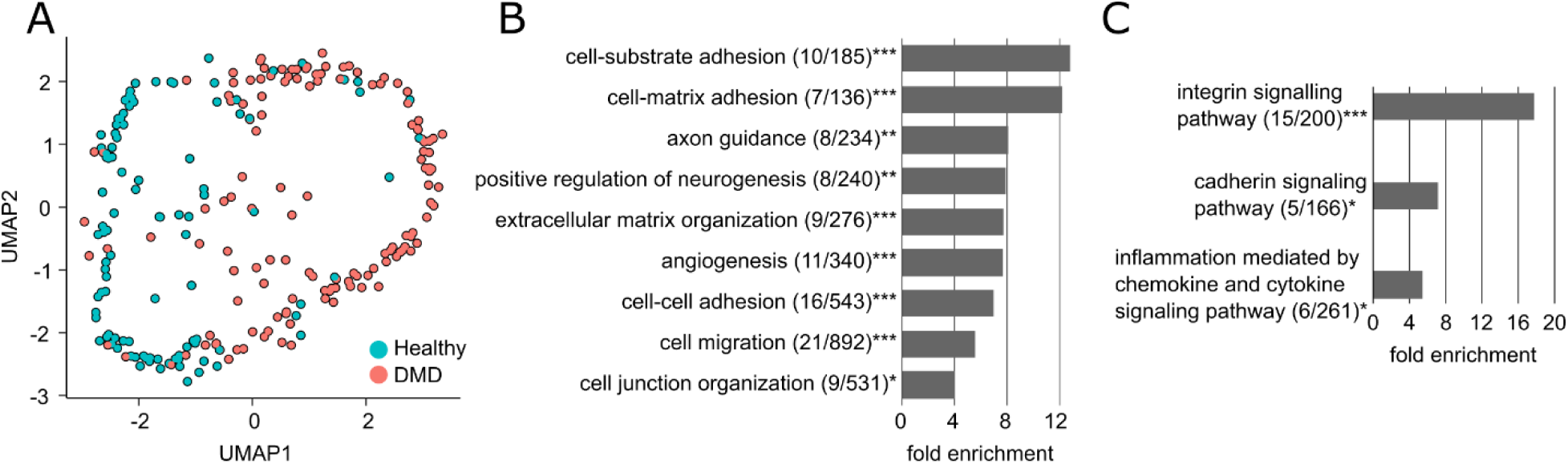
DMD cells exhibit a marked dysregulation of cell junction genes at Day 10. (A) UMAP plot of the individual cells collected at Day 10 from the sci-RNA-Seq data set (Figure 1) colored by their hiPSC line of origin. (B-C) Gene Ontology (B) and Pathway enrichment (C) analyses performed with the 94 genes differentially expressed in DMD cells at Day 10 in the sci-RNA-Seq data set. Legend: * false discovery rate (FDR) < 0.05; ** FDR < 0.01; *** FDR < 0.001; (X/Y): number of genes from the GO category found differentially expressed in the dataset / total number of genes in the GO category.

### Dystrophin deficiency does not compromise myotube formation despite upstream dysregulation of multiple cell junction protein families

The dysregulation events triggered by dystrophin deficiency in somite progenitors were evaluated using a myogenic differentiation protocol that utilizes media formulation commercialized by amsbio and previously shown to generate a homogeneous muscle cell population^17,20^. In addition to the DMD patient hiPSC line, an isogenic DMD CRISPR line was engineered from the healthy control line by deleting the entire exon 45 of the *DMD* gene together with 17 base pairs in exon 54^38^, leading to an absence of dystrophin protein (Figure S2A). The cell lines and the differentiation protocols used at each step of this study are recapitulated in Table S4. The ability of DMD patient-derived and CRISPR-engineered dystrophin null hiPSCs to differentiate into multinucleated myotubes using the amsbio protocol was evaluated by immunofluorescence after 25 days of differentiation. Myotubes were stained for two skeletal muscle proteins: myosin heavy chains and α-actinin. Multinucleated myofibers with striations could be observed in the 3 hiPSC lines (Figure 3A, Figure S2B). We quantified the area positive for α-actinin and obtained comparable fluorescent signals in the 3 hiPSC lines representing 34 % of the total area on average (Figure 3B). Thus, overall myotube formation is not impacted by dystrophin deficiency with the amsbio protocol. Prior to terminal differentiation, hiPSCs undergo several rounds of amplification and passages as they transition through the successive developmental intermediates. In particular, a “somite” gene expression signature was previously described by bulk RNA-Seq at differentiation Day 10^20^. Thus, hiPSCs were differentiated up to Day 10 and analyzed by single-cell RNA-Seq. After dimensional reduction, the UMAP plot revealed that DMD cells and healthy controls were separated into distant clusters. The CRISPR clusters were found in between, closer to the DMD cells with only a small fraction overlapping with healthy cells (Figure 3C). Expression of well-characterized somite marker genes such as *MET*, *PTN*, *NR2F1* and *NR2F2*, *MEOX1* and *PAX3* was confirmed at single-cell resolution (Figure 3D). Importantly, DMD patient cells expressed significantly higher levels of *PAX3*, *MEOX1* and *NR2F1* than healthy controls, and lower levels of *MET*, *PTN* and *NR2F2*. The CRISPR line presented with a less pronounced dysregulation profile, yet statistically significant for all the above-mentioned marker genes except *NR2F1*. Absence of differentiation by-products was confirmed at Day 10, with little to no expression of neural tube (*SOX2*, *PAX6*, *IRX3*), lateral plate mesoderm (*GATA4*), intermediate mesoderm (*PAX8*) and sclerotome (*PAX1*) marker genes on the UMAP, confirming the overall purity of the cultures (Figure S3A). Differential expression analysis using the Monocle 3 pipeline identified 4,565 and 2,913 significantly dysregulated genes in the DMD patient line and the CRISPR line, respectively, when compared to the healthy control (p-adj < 0.01, Table S5). Among these, 1,885 genes were differentially expressed in both lines, which represents the core gene set dysregulated at the somite stage as a direct consequence of the DMD deficiency (Figure 3E). Of note, this also illustrates the influence of the genetic background on disease manifestation at the transcriptome level, as 36 % of the CRISPR-induced dysregulations were not present in the DMD patient line. Importantly however, the common DEGs also included 55 of the 94 genes (59 %) found significantly dysregulated at Day 10 in the previous dataset (Figure 2). GO analysis on the 1,885 overlap genes showed enrichment for terms related to cell junctions, such as desmosome organization (p = 7.2E-3), tight junction organization (p = 4.4E-4), and cell junction assembly (p = 2.2E-11) (Figure 3F). Specific focus on key cell junction families highlighted marked down-regulations of tight junction genes such as claudins (*CLDN*), ocludin (*OCLN*) and tight junction proteins (*TJP*), but also desmosome genes such as desmoplakin (*DSP*), desmogleins (*DSG*) and desmocollins (*DSC*) in the DMD line. In contrast, multiple genes from the protocadherin (*PCDH*) and cadherin (*CDH*) families were up-regulated (Figure 3G). As above, the CRISPR line showed an intermediary dysregulation profile for all cell junction protein families (Figure 3H). In the initial dataset, adherens junctions including *CDH* and *PCDH* genes were also part of the BEAM gene list (Table S2), together with genes encoding catenins (*CTNN*) (Figure S3B). Altogether, these results suggest a major role for cell junction gene families in the initiation and progression of DMD during development.

**Figure 3:**
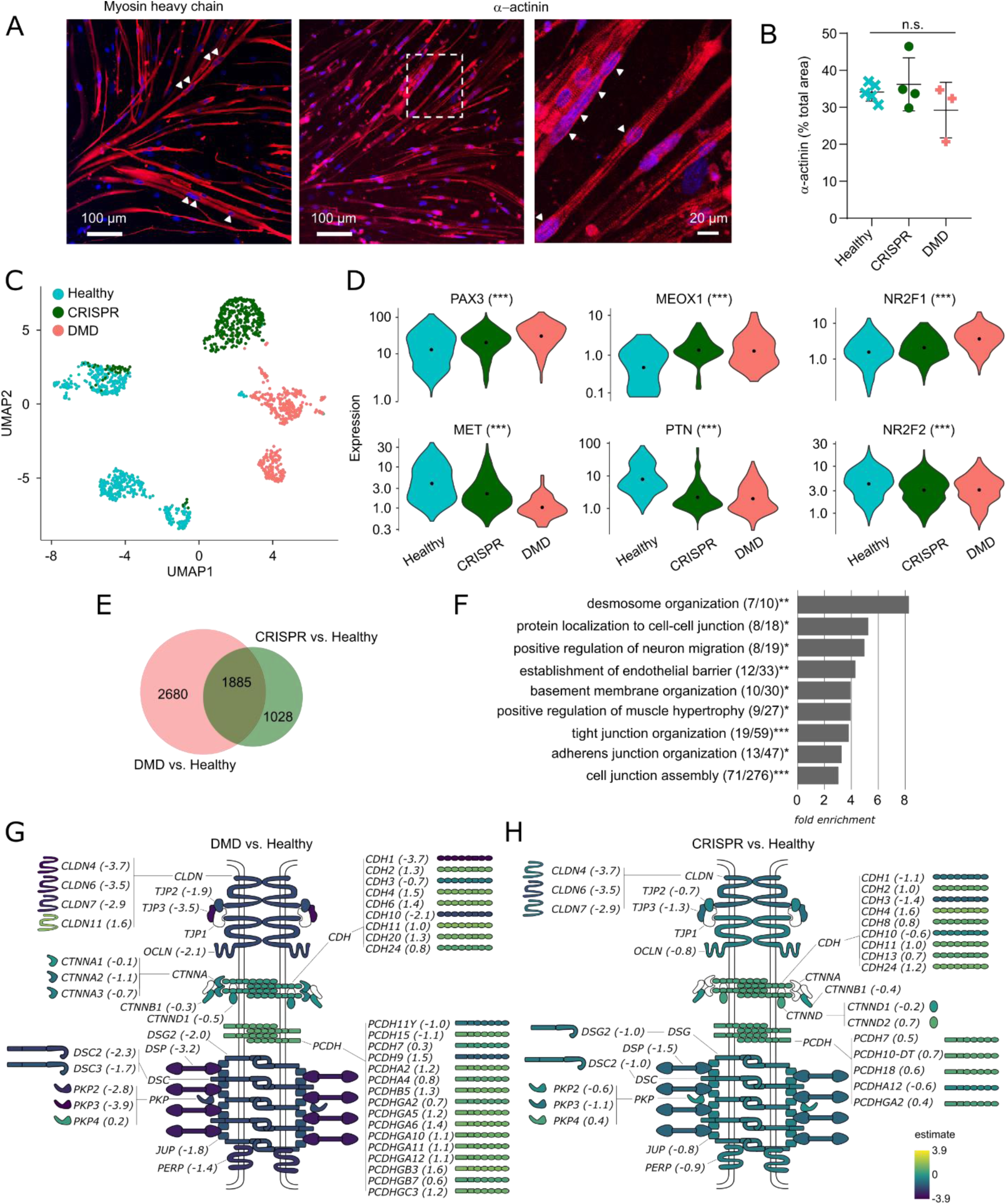
Dystrophin deficiency does not compromise myotube formation despite upstream dysregulation of multiple cell junction protein families. (A) Fluorescent staining of myosin heavy chains and a-actinin (red) in myotubes derived from Healthy hiPSCs. Arrowheads indicate multinucleation and the right panel focuses on a striation pattern. (B) Quantification of the α-actinin fluorescent area in the three cell lines, expressed as a percentage of total area (n = 3 to 5 panels by cell line). (C) UMAP plot showing the 3566 individual cells colored by their hiPSC line of origin. (D) Violin plots showing the expression of somite marker genes in single cells from Healthy, CRISPR and DMD hiPSCs at Day 10. (E) Venn diagram of the differentially expressed genes in DMD and CRISPR hiPSC lines at Day 10. The absolute numbers of genes are indicated in the appropriate sections. (F) Gene ontology analysis with the 1885 genes differentially expressed in both the DMD and the CRISPR hiPSC lines at Day 10, showing a selection of significantly enriched biological processes. Legend: * false discovery rate (FDR) < 0.05; ** FDR < 0.01; *** FDR < 0.001; (X/Y): number of genes from the GO category found differentially expressed in the dataset / total number of genes in the GO category. (G-H) Differential expression of cell junction genes and potential implications at the protein level in the DMD (G) and CRISPR (H) hiPSC lines. Fold change estimates from single-cell data are indicated between brackets and color-coded for each protein. CLDN: claudin; OCLN: occludin; TJP: tight junction protein; CDH: cadherin; CTNN: contactin; PCDH: protocadherin; DSG: desmoglein; DSP: desmoplakin; DSC: desmocollin; PKP: plakophilin; JUP: plakoglobin; PERP: p53 apoptosis effector related to PMP22.

### Dystrophin deficiency leads to impaired cell state transitions during *in vitro* somite development

The role of cell junction proteins in the dynamics of paraxial mesoderm development have been highlighted in multiple species, particularly in the cell state transitions happening during the formation of somites bilaterally on either side of the neural tube and later during the delamination of the dermomyotome^10–12,39–46^. Thus, hiPSC-derived somite progenitors *in vitro* might undergo successive mesenchymal-to-epithelial and epithelial-to-mesenchymal transitions accompanied by a well-coordinated remodeling of the cell-cell and cell-matrix junctions. The morphology of the hiPSC cultures was evaluated between Day 7 and Day 10 of the amsbio protocol. At Day 10, two cell populations with distinct features were observed: (1) spindly, refringent, tightly packed cells with a mesenchymal morphology and (2) larger, more flattened cells in close contact to one another similar to squamous or cuboidal epithelia (Figure 4A). These larger cells were first detected at Day 8 and progressively form islets until Day 10 (Figure S4A). Immunofluorescence assays showed that the islets expressed high levels of E-cadherin (E-cad), a cell junction protein characteristic of epithelial cells and were negative for Vimentin (Vim), a cytoskeleton protein expressed mostly in mesenchymal cells (Figure 4B, Figure S4B). Conversely, the cells surrounding the islets expressed Vim but no E-cad. The fluorescent signal was quantified in the 3 hiPSC lines at Day 10 and we found ∼10% E-cad-positive cells in both the Healthy control line and the CRISPR line, and less than 1% in the DMD line (Figure 4C). We next assessed the expression of C-Met, a receptor tyrosine kinase expressed at the membrane of epithelial cells during dermomyotome development from the somites and critical for subsequent myoblast migration^47–49^. Interestingly, we observed a strong C-Met expression in epithelial islets, suggesting a dermomyotome identity, but no C-Met-positive cell was found in the DMD line. Differentiation of the CRISPR line resulted in C-Met expression at Day 10, but at a level 5 times lower than in the healthy control (p-val = 0.029) (Figure 4D-E). This suggests that somitogenesis and the associated cell state transitions are altered in the absence of dystrophin, but that additional factors have an influence, depending on the genetic background.

**Figure 4:**
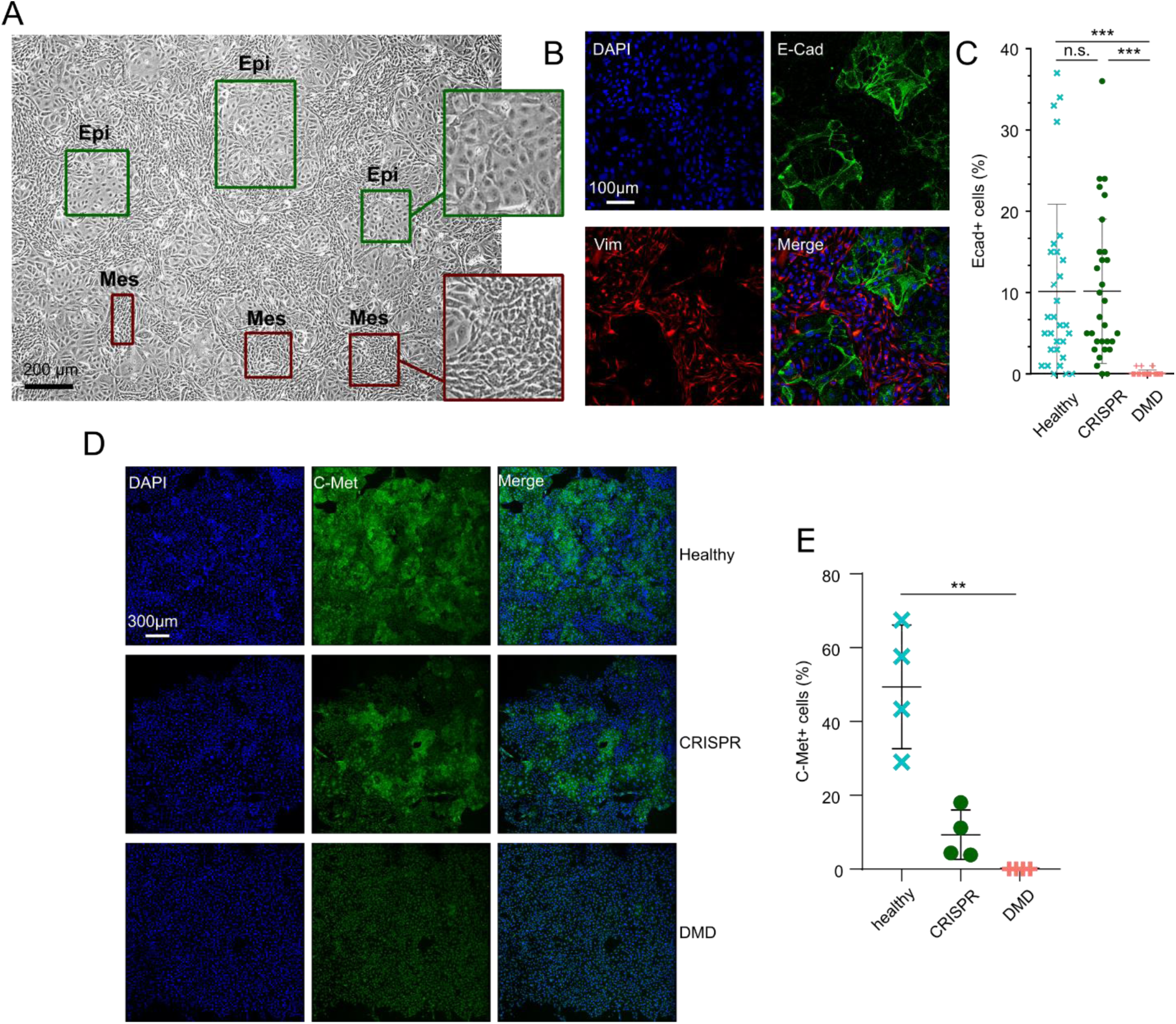
Dystrophin deficiency leads to impaired cell state transitions during in vitro somite development. (A) Optical microscopy with phase contrast of somite progenitors derived from hiPSCs. Insets highlight the “epithelial-like” (Epi) and “mesenchymal-like” (Mes) cell populations. Scale bar = 200µm. (B) Detection of E-Cad and Vim in somite progenitors derived from hiPSCs by immunofluorescence and confocal microscopy. (C) Quantification of the percentage of E-Cad-positive cells in the Healthy, DMD and CRISPR hiPSC lines at Day 10 in three replicate experiments (N = 10 panels by experiment). n.s. not significant; *** p-value < 0.001. (D) Detection of C-Met in the three hiPSC lines by immunostaining and confocal microscopy at differentiation Day 10. (E) Quantification of the C-Met fluorescent area normalized by the number of nuclei. Imaging was performed on four large mosaic panels per cell line.

### DMD-specific defects in somitogenesis and cell junction gene dysregulations are independent of the genetic background

To confirm that cell junction dysregulation during embryonic myogenesis in the absence of dystrophin was also observed in other genetic backgrounds, we compared our current findings with an RNA-Seq dataset previously generated by our group that describes myogenic differentiation of DMD hiPSCs at defined time points (https://muscle-dmd.omics.ovh/)^20^. More precisely, hiPSC lines from three independent DMD patients and three healthy individuals were differentiated in triplicate with the amsbio media formulation^17^ and cells were collected for bulk RNA-Seq at several time points (Figure S5A). Interestingly, the transcriptome of DMD hiPSCs markedly diverged from healthy controls at Day 10, when somite marker genes are expressed^20^. Immunostaining at Day 10 showed the presence of epithelial islets expressing C-Met in the 6 hiPSC lines, but at a level 5.7 lower in the 3 DMD patient lines, confirming the results obtained in the initial DMD patient-derived and CRISPR lines (Figure 5A-B). Further analysis of the RNA-Seq data set identified 1450 genes significantly dysregulated at Day 10 in the 3 DMD cell lines (abs(log2 fold-change) ≥ 1 and adjusted p-value ≤ 0.01), among which several cell junction protein families (Figure 5C, Table S6). GO analyses showed a significant enrichment in genes involved in the formation of cell junctions (Figure 5D), which confirms their importance in the initial manifestation of DMD during somite differentiation, independently of the genetic background. Particularly, desmosome organization (GO:0002934) was affected in DMD cells, with a significant downregulation of *GRHL1* (log FC = -2.5, p-val = 3.7E-06), *JUP* (log FC = -1.0, p-val = 7.7E-04), *PERP* (log FC = -1.4, p-val = 1.2E-08), *PKP2* (log FC = -1.5, p-val = 2.6E-03) and *PKP3* (log FC = -2.9, p-val = 1.1E-03) (Figure 5E). Tight junction organization (GO:0120193) was also perturbed by the absence of dystrophin, with dysregulation of gene members of the claudin (CLDN), cadherin (CDH) and protocadherin (PCDH) families (Figure 5E). Overall, cell junction genes were mostly downregulated in DMD cells, with the exception of the protocadherin (*PCDH*) family, in which 8/9 genes were upregulated (*PCDH17*, *PCDHA2*, *PCDHA7*, *PCDHB18P*, *PCDHGA3*, *PCDHGA4*, *PCDHGA5*, *PCDHGB3*, mean log FC = 1.5) and only 1 was downregulated (*PCDHA13*, log FC = -1.7). To evaluate the functional impact of cell junction gene dysregulation at Day 10, the migration velocity of DMD somite progenitors was compared to healthy control cells with a scratch-wound assay. Cell migration was precisely quantified by the relative wound density (RWD), which measures the cell density in the initial wound area normalized by the density outside the wound area of the same well. Healthy control cells were shown to reach a RWD of 50% in 7.8 +/- 0.5 hrs, and the wound area was almost entirely closed in the first 12 hrs (Figure 5F). Given this time frame, the impact of cell division was assumed minimal compared to the intrinsic cell migration. Temporal follow-up of wound area recolonization showed that DMD cells migrated 17 % faster than healthy controls with RWD50 = 6.5 +/- 0.1 hrs (p = 0.03), in agreement with the defective epithelialization and the predominance of a mesenchymal population (Figure 5G-H). Similarly, the time taken by DMD cells to restore a confluency of 50% in the wound area was 34% lower in DMD cells (6.5 +/- 0.1 hrs *vs.* 9.5 +/- 0.4 hrs in Healthy controls) (Figure S5B). These results strongly suggest that dystrophin deficiency has functional consequences during myogenic development, particularly when somite progenitors undergo successive cell state transitions resulting in delamination and colonization of future muscle territories.

**Figure 5:**
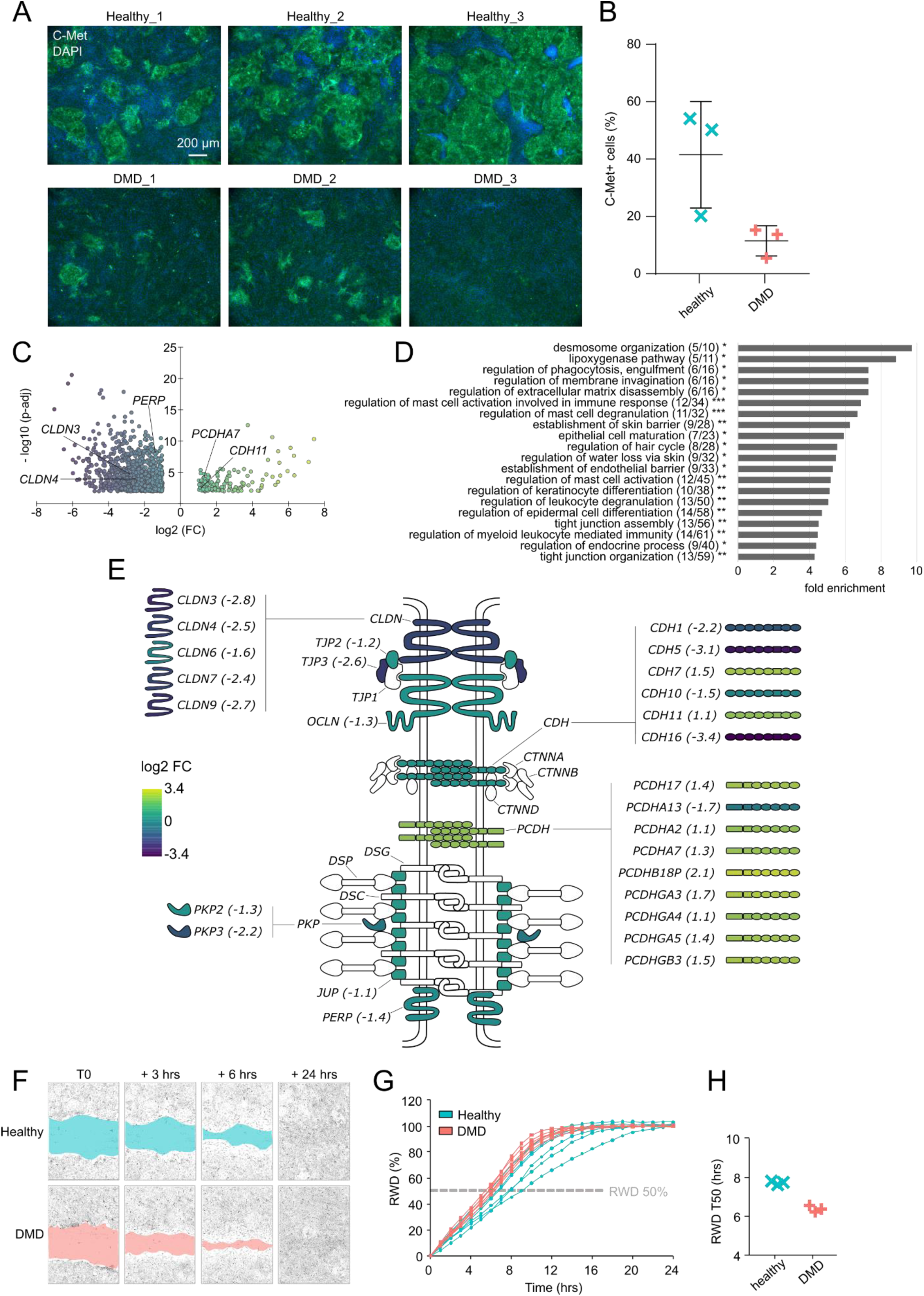
DMD-specific defects in somitogenesis and cell junction gene dysregulations are independent of the genetic background. (A) Fluorescent staining of C-Met in DMD hiPSCs and healthy control lines at Day 10 of the myogenic differentiation. One representative picture is shown per individual line (Healthy_1 to 3 and DMD_1 to 3). (B) Quantification of the C-Met fluorescent area normalized by the number of nuclei. Imaging was performed on three independent pictures per line and the data shows the median value for each line. (C) Volcano plot of the differentially expressed genes in DMD somite progenitor cells at Day 10 (thresholds: |log2 FC| > 1 & p-adj < 0.01). (D) Gene Ontology analysis using the differentially expressed genes as input and showing key biological processes. Legend: * false discovery rate (FDR) < 0.05; ** FDR < 0.01; *** FDR < 0.001; (X/Y): number of genes from the GO category found differentially expressed in the dataset / total number of genes in the GO category. (E) Differential expression of cell junction genes and potential implications at the protein level. Fold change values (DMD vs. Healthy) are indicated between brackets and color-coded for each protein. CLDN: claudin; OCLN: occludin; TJP: tight junction protein; CDH: cadherin; CTNN: contactin; PCDH: protocadherin; DSG: desmoglein; DSP: desmoplakin; DSC: desmocollin; PKP: plakophilin; JUP: plakoglobin; PERP: p53 apoptosis effector related to PMP22. (F) Representative scratch wound assay in Healthy and DMD cells at Day 10 (T0). Recolonization of the wound area is shown by the reduction of the blue/red area. (G) Monitoring of the relative wound density (RWD) over time in the 3 DMD and 3 Healthy control lines. (H) For each line, the time to reach an RWD of 50% (RWD T50) was determined and averaged from 2 x 8 wells in 2 independent experiments.

Altogether, our study combines data from four unrelated DMD patient hiPSC lines and four healthy controls plus one isogenic DMD mutant, and indicates that dystrophin deficiency leads to major dysregulations of cell junction gene families participating in cell state transitions during somite development. As a working model, dysregulated somitogenesis and aberrant cell migration may result in the developmental phenotypes previously observed in foetuses at risk of developing DMD (Figure 6).

**Figure 6:**
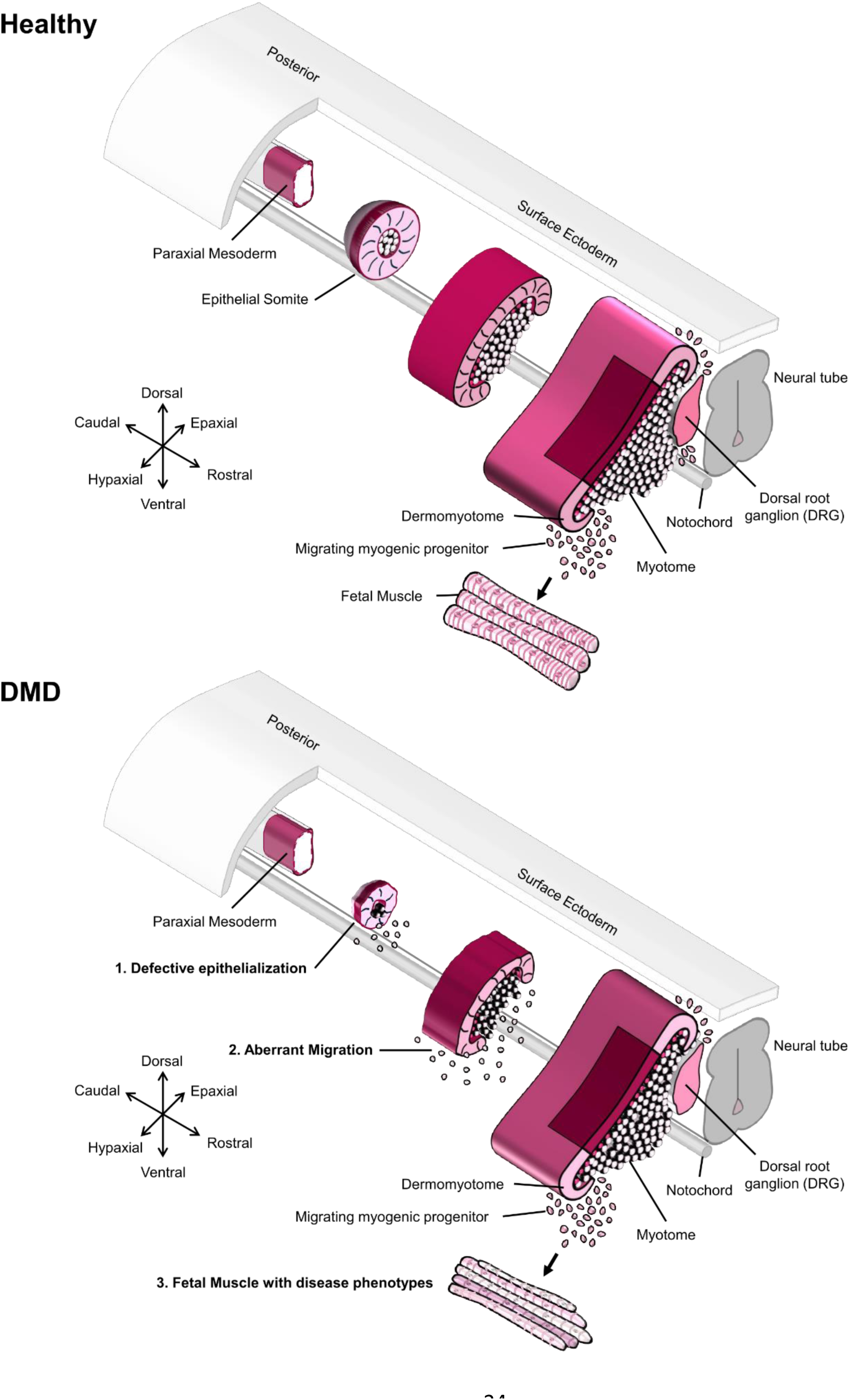
Working model of dysregulated somite development in DMD embryos. The top panel shows the successive steps of somitogenesis from the paraxial mesoderm in a healthy context, including the migration of myogenic progenitors colonizing the fetal muscle territories. In the bottom panel, the defective epithelialization and the aberrant migration phenotypes are highlighted in DMD somites, giving rise to fetal muscles with disease phenotypes, as previously published^7,8^.

## Discussion

Thanks to carefully defined differentiation protocols, hiPSCs and their derivatives can help better understand human embryonic development and investigate the early impact of mutations leading to specific genetic disorders without the use of embryos^14^. Using a published protocol recapitulating key developmental steps of human myogenesis, we provide here a temporally resolved dataset spanning ten time points along the differentiation of healthy and DMD hiPSCs from pluripotency to the skeletal muscle lineage. Pluripotent cells first go through intensive transcriptional changes as they leave pluripotency and adopt an early mesodermal fate. After Day 7, they stabilize as they become somite progenitors and their progenies (*i.e.* dermomyotome and skeletal muscle cells). The developmental trajectory computed from pseudotime data showed that fractions of cells reach the end of the myogenic process as early as Day 10, reflecting the fact that differentiation is heterogeneous and asynchronous^33,50,51^. During the second half of the differentiation process, cells that are “behind schedule” probably “catch up” or remain stalled on an abortive developmental path. At the differentiation end stage, a majority of cells did not express canonical myogenic markers such as *MYOD1*, *MYOG* or *PAX7*. However, most of them were positive for the myocyte-specific enhancer factor 2C (*MEF2C*), known to be involved in myogenesis but also in the development of a myriad of other tissues (*i.e.* heart, nervous system, vasculature, bone and cartilage)^52^. Neural cells have already been identified as cellular by-products generated alongside muscle cells with the myogenic differentiation protocol used in the first part of this study^37^. For this reason, a commercial myogenic media known to produce more homogeneous cell populations at the successive developmental steps was used in the subsequent steps of this study^17,20,53^ (Table S4). This allowed us to confirm that similar to what is seen in DMD patients, the absence of dystrophin does not compromise the ability of hiPSCs to differentiate into myotubes. However, it may delay differentiation and / or deflect a fraction of cells from the skeletal muscle into alternative or abortive lineages during embryogenesis, which might go unnoticed *in vitro* due to the amplification and passage steps that are part of the differentiation protocols.

Here, we discovered that dystrophin mutation leads to the dysregulation of cell junctions during the successive cell state transitions which come with the *in vitro* differentiation of “somite-like” cell monolayers. Somites are transient metameric structures emerging from the paraxial mesoderm in the embryo. They give rise to multiple tissues such as the axial dermis, cartilages, bones and skeletal muscles of the axis and limbs^54,55^. Individual somites are patterned along the dorsal-ventral axis into the sclerotome at the ventral pole and the dermomyotome at the dorsal pole. Myogenic progenitors delaminate from the dermomyotome to form primary multinucleated myofibers^12,56,57^ (Figure 6). These developmental steps involve coordinated gene expression and successive transitions between a mesenchymal and an epithelial state^10–12^. We showed that the hiPSC model recapitulates key features of somitogenesis when subjected to myogenic differentiation, both at the level of their morphology and gene expression program. As a monolayer, somite progenitors derived from control hiPSCs spontaneously organize into discrete “epithelial islets”, surrounded by mesenchymal cells. In the embryo, somite epithelialization requires the activity of *Paraxis* and the somites of *Paraxis-/-* murine embryos showed dysregulated somitogenesis and disorganized cell junctions^11,58^. The activity of the Rho family small GTPases Cdc42 and Rac1 also plays an essential role in the epithelialization during somitogenesis^59^, possibly by controlling the architecture of the cytoskeleton and the remodeling of cell junctions during the successive cell state transitions. Besides, cell junction proteins were massively associated with somite formation, segmentation and specification in multiple species. For instance, cadherin 1, 2, 4, 5, 11 and 23 together with protocadherin 1, 8 and 18 were found expressed in developing somites of chicken embryos^40^. In the zebrafish, knockdown of cadherin 1 gene expression with morpholinos induced an aberrant somite morphology phenotype^41^. During somite segmentation in *Xenopus*, the activity of protocadherin 8 / PAPC is required for segmentation^42^, and dominant negative forms of type I cadherins impaired the rotation of segmented somitomeres^43^. The role of PAPC in somite segmentation was later confirmed in chicken embryos, where it affects the epithelialization of mesoderm cells^44^. In the mouse embryo, N-cadherin and cadherin 11 were found expressed during the early steps of somite condensation from the paraxial mesoderm and then restricted to the dermomyotome and the sclerotome, respectively^45^. In addition, the removal of claudin 3, 4 and 8 led to irregularly shaped, fused somites and defects in neural tube closure^46^. In this study, dystrophin deficiency in somite progenitors was linked with dysregulation of multiple cell junction genes including *CDH1*, *CDH2*, *CDH11*, *CLDN3*, *CLDN4*, defective epithelialization and increased migration (Figure 6). One appealing hypothesis would be that during somite development, the embryonic isoform of dystrophin serves as a stabilizing anchor for cell junction proteins in newly formed epithelial cells. Testing this hypothesis will require further investigations in appropriate model systems, as Dp412e is specific to a subgroup of anthropoids, including humans, chimpanzees, gorillas and orangutans^21^. Using our hiPSC model, we demonstrated that as a result of defective epithelialization, DMD progenitors have increased migration properties at the somite stage, which could in turn dysregulate the delamination process and the colonization of future skeletal muscle territories.

Multiple protein isoforms have been described from the *DMD* locus, each with a defined expression pattern^60–62^, but their function remains to be precisely characterized. In skeletal and cardiac muscles, the Dp427m dystrophin isoform is known to interact with the dystrophin-associated protein complex (DAPC), but in other tissues, the protein partners of other isoforms are less characterized. Recent studies have described association of dystrophin with alternative partners in specific cells and tissues, such as aquaporin 4, calcium and potassium channels^63–67^. However, the exact molecular mechanisms driven by these interactions reman to be investigated. Dp412e was identified as an “embryonic” isoform induced during the formation of early mesoderm progenitors and embryoid bodies from hiPSCs^21^. The results of this study strongly suggest that Dp412e has specific roles during embryonic development and the cell transitions occurring when somites develop. In DMD patients, the absence of this isoform during the embryonic period might trigger a cascade of subtle molecular events leading to cellular phenotypes which only become apparent when the muscle tissue reaches a certain level of maturation. Reproducing *in vitro* the conditions in which skeletal muscle normally develop after birth will be critical to further characterize the dynamics of DMD using hiPSCs.

Altogether, our study leads us to consider DMD even more as a developmental disease, even though patients are born without apparent symptoms. A better understanding of the “invisible” DMD initiation events during development and early postnatal life will help identify biomarkers early in the course of disease progression, which in turn could accelerate diagnosis and pinpoint new therapeutic targets.

## Acknowledgments

This work was funded by the INSERM ATIP Avenir program, Nantes Université, the University Hospital of Nantes, the Association Française contre les Myopathies (AFM) Téléthon, Genopole, and by the Wellstone Muscular Dystrophy Cooperative Research Center supported by the National Institute of Health (NIH, grant number U54 AR065139). We thank the GenoA Genomics Core Facility and the BiRD Bioinformatics Core Facility (Nantes Université, SFR Bonamy, UMS Biocore, Biogenouest), which is part of the Institut Français de Bioinformatique (ANR-11-INBS-0013) for the access to the computing and storage infrastructure and their help with sequencing and processing the single-cell libraries. We acknowledge the IBISA MicroPICell facility (Nantes Université, SFR Bonamy, UMS Biocore, Biogenouest), member of the national infrastructure France-Bioimaging supported by the French national research agency (ANR-10-INBS-04), for their help with fluorescence image acquisition and analysis.

## Author contributions

Conceptualization: CP, DLM, JBD; Methodology: EMo, CL, JBD; Software: VM, JSP, CT, JBD; Validation: EMo, CL, JBD; Formal analysis: EMo, CL, JBD; Investigation: EMo, EMa, VM, CL, AB, DLJ, JBD; Resources: AB, CT, OA, CP, DLM, JBD; Data Curation: EMo, VM, JSP, JBD; Writing – Original Draft: EMo, CL, JBD; Writing – Review & Editing: EMo, EMa, VM, CL, CT, CLG, OA, CP, DLM, JBD; Visualization: EMo, CL, JS, JBD; Supervision: CT, CLG, OA, CP, DLM, JBD; Project administration: JBD; Funding acquisition: EMa, VM, OA, CP, DLM, JBD.

## Inclusion and diversity

Our research group supports inclusive, diverse and equitable conduct of research.

## STAR Methods

### RESOURCE AVAILABILITY

#### Lead contact

Further information and requests for resources and reagents should be directly addressed to and will be fulfilled by the lead contact, Jean-Baptiste Dupont.

#### Materials availability

This study did not generate new unique reagents

#### Data and code availability

- The two single-cell RNA-seq data sets have been deposited at GEO and are publicly available as of the date of publication. Accession numbers are listed in the key resources table. This paper also analyzes publicly available data. These accession numbers are listed in the key resources table. Original western blot images and microscopy data reported in this paper have been deposited at Zenodo repository. DOIs are listed in the key resources table.
- All original code has been deposited as R Notebook files at Zenodo Repository. DOIs are listed in the key resources table.
- Any additional information required to reanalyze the data reported in this paper is available from the lead contact upon request.

### EXPERIMENTAL MODEL AND STUDY PARTICIPANT DETAILS

#### Ethical statement

The Healthy, DMD and CRISPR lines have been described, characterized and published previously^20,35,38^. Participants gave informed consent for the generation of urine-derived hiPSC lines, as required by the Institutional Review Board (IRB). The experiments performed in this manuscript fall into the Category 1A described in the Guidelines for Stem Cell Research and Clinical Translation of the International Society for Stem Cell Research (ISSCR, https://www.isscr.org/guidelines).

#### hiPSC lines

The Healthy UC3-4 line was generated from an adult male participant with no known skeletal muscle disease. The DMD 72039 line was generated from an adult male with a declared DMD pathology and carrying an exon50 deletion in the *DMD* gene. These individuals provided urine samples, from which urine cells were expanded and reprogrammed to generate hiPSCs by Guan X. *et al.* at the University of Washington, Seattle (USA)^35^. The DMD CRISPR line was generated from the UC3-4 line by CRISPR Cas 9 gene editing by Smith A. *et al.* in collaboration with the Institute for Stem Cell and Regenerative Medicine Ellison Stem Cell Core^38^. It carries an exon 45 deletion and a 17 bp deletion in exon 54 of the *DMD* gene. The absence of dystrophin was confirmed by Western blot after myogenic differentiation, in comparison with the Healthy UC3-4 control line (Figure S2A). Cell lines were not otherwise authenticated. The three hiPSC lines were extensively characterized, expanded and shared with the TaRGeT INSERM laboratory at Nantes Université through a dedicated Material Transfer Agreement. The three additional DMD patient-derived hiPSC lines and the three additional healthy control lines were previously generated by Mournetas V. and Massouridès E. at the I-Stem institute, Corbeil-Essonnes (France)^20^. The mutations are specified in Figure S4.

#### hiPSC maintenance

All cell culture experiments were performed at 37 °C and 5 % CO2 in a standard tissue culture incubator. The three hiPSC lines were expanded in the TaRGeT laboratory as Master Cell Banks (MCB) at various passages (Healthy: passage 34; DMD: passage 25; CRISPR: passage 46). Cells were thawed at 37°C and seeded as clusters on Matrigel-coated plates (1:60 dilution) after spinning 3 min at 300 g and resuspension in mTeSR Plus culture medium supplemented in 10 µM ROCK inhibitor. Fresh media without ROCK inhibitor was renewed the day after seeding and then every other day. Cultures were manually cleaned from abnormally looking clusters and were passaged with Versene when the overall confluence reached 70 – 80 %. Working Cell Banks (WCB) were stored after two passages from the MCBs. Cells were detached with Versene, spun down 3 min at 300 g, and clusters were gently resuspended in Cryostor freezing media for long term cryopreservation in liquid nitrogen.

#### Rat Muscle biopsies

The protein samples used as positive controls in the dystrophin western-blot were obtained from rat pectoral muscle biopsies from a previous study^68^. Dmd*^mdx^* rats and healthy controls were handled and housed in the UTE IRS2 from Nantes Université, according to a protocol approved by the Institutional Animal Care and Use Committee of the Région des Pays de la Loire (University of Angers, France) as well as the French Ministry for National Education, Higher Education and Research (authorization #2018102616384887).

### METHOD DETAILS

#### hiPSC myogenic differentiation

This study combines the use of two myogenic differentiation protocols:

- Establishment of the myogenic trajectory: hiPSCs were differentiated with a succession of five defined media, as published previously^16^. After seeding on Matrigel (1:60 dilution) as single cells at a density of 30,000 cells / cm², hiPSCs were grown for 2 – 3 days in mTeSR Plus media. ROCK inhibitor was added upon seeding but removed by a fresh media change after one day. The cells were then incubated in five successive differentiation media based on Dulbecco’s Modified Eagle’s Medium (DMEM) / F12 supplemented with Non-Essential Amino Acids (NEAA) and additional molecules detailed below:

`o` Media 1 (3 days): Insulin transferrin selenium (ITS) 1X + 3 µM CHIR99021 + 0.5 µM LDN
`o` Media 2 (3 days): Insulin transferrin selenium (ITS) 1X + 3 µM CHIR99021 + 0.5 µM LDN + basic Fibroblast Growth Factor (bFGF) at 20 µg / ml
`o` Media 3 (2 days): 15 % Knockout serum replacement (KSR) + 0.5 µM LDN + bFGF at 20 µg / ml + Hepatocyte Growth Factor (HGF) at 10 µg / ml + Insulin-like Growth Factor (IGF) 1 at 2 ng / ml.
`o` Media 4 (4 days): 15 % KSR + IGF-1 at 2 ng / ml
`o` Media 5 (16 days): 15 % KSR + IGF-1 at 2 ng / ml + HGF at 10 µg / ml

Cells were collected at Day 0, 1, 2, 4, 7, 10, 14, 18, 22 and 28 after incubation with Trypsin-EDTA alone, or with a combination of Trypsin-EDTA + collagenase IV at 50 U / µl for 10 min at 37 °C, mechanical dissociation and passage through a 70 µm cell strainer to remove debris and extracellular matrix.

- Post hoc analysis of the somite differentiation stage at Day 10: in parallel, hiPSCs were differentiated with a commercially available media previously shown as capable of reproducing the somite stage with high accuracy^17,20^. This protocol uses lower initial seeding densities and allows for an easier visualization and imaging of the cultures as differentiation progresses. Briefly, hiPSCs were first amplified and dissociated as single cells as described above, and then seeded on collagen I-coated plates at 3,500 cells / cm² in Skeletal Muscle Induction Medium (amsbio SKM01). The media was changed every 2 to 3 days. At Day 7, cells were dissociated with Trypsin-EDTA and cryopreserved in CryoStor CS10 medium. They were seeded on new collagen-coated plates at 20,000 cells / cm² in SKM01 until Day 10.

#### single-cell RNA-Seq

- sci-RNA-Seq: the myogenic trajectory followed by hiPSCs was determined by single-cell combinatorial indexing RNA-Seq (sci-RNA-Seq) as previously published^36^. Cells were collected as differentiation progressed (5 to 10 million per sample, from Day 0 to Day 28) and spiked with 10 % murine cells of the NIH/3t3 cell line for estimation of doublet proportions. after centrifugation for 5 min at 300 g and 4 °C, pellets were washed in DPBS and resuspended in pre-chilled methanol for fixation and permeabilization. They were then stored at – 20°C until all samples were processed at the end of the differentiation. Fixed cells were pelleted by centrifugation (same settings), washed twice in 1 ml DPBS + 1 % Diethylpyrocarbonate (DEPC), and another three times in 1 ml of cell wash buffer containing 1 % SUPERase In RNase Inhibitor and 1 % BSA in ice-cold DPBS. The final resuspension was made in 100 µl of cell wash buffer before counting with a hemocytometer. The Reverse Transcription step was performed 10 min at 55 °C *in situ* on 2,000 cells per sample, with the Superscript IV RT kit and barcoded oligo dT primers dispatched in a Lo-bind 96-well plate. To increase the diversity of barcode combinations obtained after indexing, six distinct barcodes were used for the Day 0, Day 1 and Day 2 samples and four barcodes for the remaining samples (Day 4, Day 7, Day 10, Day 14, Day 18, Day 22, Day 28). The RT reaction was stopped with 40 mM EDTA and 1 mM spermidine (5 µl per well). Cells with barcoded cDNA were then pooled in a flow cytometry tube, stained with 300 µM DAPI and sorted in a new 96-well Lo-bind plate (25 cells per well) containing Elution buffer (5 µl per well). Sorting plates can be stored at – 80 °C after brief centrifugation. Second strand synthesis was performed in each well with the NEBNext® Ultra™ II Non-Directional RNA Second Strand Synthesis Module (0.5 µl of Buffer + 0.25 µl of enzyme per well). After incubation at 16 °C for 150 min, the reaction was terminated at 75 °C for 20 min. Tagmentation was performed in each well with the illumina Tagment DNA TDE1 Enzyme (0.5 µl per well) and buffer (5 µl per well) kit, after addition of human genomic DNA (0.25 µl per well). The plate was incubated at 55 °C for 5 min and the reaction was stopped with 12 µl DNA binding buffer per well and incubation at room temperature (RT) for 5 min. AMPure XP beads (36 µl per well) were added for purification with standard protocol. The elution step was performed in a final volume of 17 µl. Libraries were amplified by PCR using the NEBNext High-Fidelity 2X PCR Master Mix and barcoded P5 primers (2 µl of 10 µM primer per well), in addition to a 10 µM P7 primer (2 µl per well). Amplification was carried out in a standard thermal cycler with the following program: 5 min at 72 °C + 30 sec at 98 °C + 18 cycles of (10 sec at 98 °C + 30 sec at 66 °C + 30 sec at 72 °C) + 5 min at 72 °C. Samples were collected from each well and pooled in a single tube. The library was then purified using 0.8 volume of AMPure XP beads according to manufacturer’s instructions. Libraries were quantified by Qubit and visualized by electrophoresis on a 6 % TBE-PAGE gel. Sequencing was performed on the NextSeq 500 platform using a V2 75 cycles kit and the following settings: Read 1: 18 cycles, Read 2: 52 cycles, Index 1: 10 cycles, Index 2: 10 cycles. Here, two sorting plates were generated and used to generated independent libraries sequenced successively with distinct P7 primers. The first sorting plate was full and thus 96 barcoded P5 primers were used in the PCR amplification step; the second sorting plate only used 86 wells (and thus 86 barcoded P5 primers). The sequences of the barcoded primers used in this protocol can be found in Table S7. Raw data analysis involved base calling (bcl2fastq), demultiplexing based on P5 barcodes (1 mismatched base allowed), adaptor trimming (trim_galore), alignment to the human (hg19) or the mouse (mm10) genome (STAR), removing of UMI duplicates and demultiplexing based on RT barcodes. Percentages of reads mapping uniquely to the human and the mouse genome were quantified and cells with over 90 % of reads assigned to the human genome were kept for subsequent analysis. Secondary analysis of the cell data set object was performed with the Monocle 3 analysis pipeline (Trapnell 2014, Qiu 2017), whose code is freely available on the Cole Trapnell lab Github and in the Zenodo Repository with the DOI listed in the key resources table.
- split-pool barcoding: the *post hoc* analysis of the somite differentiation step at Day 10 was performed with the split-pool barcoding kit commercialized by parse Biosciences, according to manufacturer’s instructions. Cells were collected at Day 10 and counted with a hemocytometer to isolate 500,000 cells for subsequent fixation and permeabilization with the Cell Fixation Kit. Samples were stored at – 80°C and thawed immediately prior to library preparation with the Evercode Whole Transcriptome Mini Kit. Primary analysis of the raw data including quality control, alignment to the human genome (hg19), demultiplexing and generation of the matrix, feature annotation and cell annotation files were carried out by the proprietary Parse Biosciences analysis suite. Secondary analysis used the Monocle 3 pipeline as previously indicated. The code has been deposited at Zenodo Repository with the DOI listed in the key resources table.

#### Immunofluorescence

- Somite progenitors: hiPSCs were first differentiated into somite progenitor cells with the amsbio commercial protocol to constitute a working cell bank at Day 7. Subsequently, cryotubes containing one million progenitors were thawed and seeded at 23,500 cells/cm² on Lab-Tek™ 4-well chamber slides (Thermofisher Cat# 154526PK) coated with collagen I (1:60 dilution). Cells were maintained in Skeletal Muscle Induction medium supplemented with 2% Pen/Strep and incubated at 37°C, 5% CO2 until confluency.
- Myotubes: myogenic progenitors were differentiated from hiSPCs with the amsbio commercial protocol and banks were made at Day 17. Subsequently, cryotubes containing one million progenitors were thawed and seeded at 23,500 cells/cm² on Lab-Tek™ 4-well chamber slides (Thermofisher Cat# 154526PK) coated with collagen I (1:60 dilution). They were maintained in Skeletal Muscle Myoblast medium supplemented with 2% Pen/Strep and incubated at 37°C, 5% CO2 until confluency. Differentiation was induced with the Skeletal Muscle Myotube medium supplemented with 2% Pen/Strep for 8 days, with a fresh media change every 2 to 3 days.

Cells were fixed in 4 % paraformaldehyde (PFA) for 1 hr and then permeabilized in PBS + Triton 1X + 2.5% bovine serum albumin (BSA). Primay antibodies were diluted in permeabilization buffer and incubated on the cells overnight at 4 °C (E-Cadherin Alexa Fluor™ 488, Vimentin eFluor™ 570 and C-Met: 1:50 dilution; Myosin Heavy Chain: 1:300 dilution; α-actinin: 1:500 dilution). The next day, nuclei were stained with 1:10.000 DAPI and for myotubes, the anti-mouse or anti-goat (C-Met) secondary antibodies were diluted at 1:1000 in PBS and added for 1 hr at RT. Coverslips were mounted on the slides using ProLong Gold Antifade Reagent after removing the Lab-Tek™ walls. Images were taken with a 20 X oil immersion objective on a NIKON® A1 RSi confocal microscope.

#### Western-blot

Proteins were extracted from frozen cell pellets in RIPA buffer during 1 hour (10 mM Tris + 150 Mm NaCl + 1mM protease inhibitor cocktails + 1% Igepal + 0.1% SDS). Protein extracts diluted at 1:10 were quantified with the DC Protein Assay kit at 750 nm on the Thermo Scientific™ Multiskan™ GO. Protein samples (50 µg per sample) were denatured by addition of NuPAGE LDS sample buffer 4X and Dithiothreitol (DTT), and loaded onto a Nupage 3-8% TA Gel before migrating for 2 hrs at 100 V in 1X NuPAGE tris acetate SDS buffer. Protein extracts from Healthy and DMD*^mdx^* rat pectoral muscles were loaded as controls. The Bio-Rad Trans-Blot Turbo Transfer System was used for protein transfer. After overnight saturation at 4 °C in saturation buffer (5 % milk + 0.1 % Tween 20 + 1 % NP40), membranes were incubated with primary Mouse anti-dystrophin NCL-DYS2 antibody diluted at 1:250 for 1 hr. The α-tubulin protein was labeled with a Mouse Monoclonal Anti-α-Tubulin antibody diluted at 1:10000. All membranes were then washed three times for 5 minutes in PBS + 0.1 % Tween 20 before incubation with the corresponding secondary antibodies (Goat Anti-Mouse antibody/HRP diluted at 1 :5000 during 1 hr). The ECL kit was used for detection of the HRP enzyme activity, with exposure times of 2 minutes (α-tubulin) and 2 hours (Dystrophin).

#### Cell migration assay

Somite progenitors derived from hiPSCs were frozen and amplified at Day 7 of the amsbio SkM differentiation protocol. They were seeded in an Imagelock 96-well plate at 20,000 cells/cm² and grown in SkM01 medium for 3 days (8 wells per line). At Day 10, the Incucyte® Woundmaker™ was used to create homogeneous wound in the cell monolayer, according to the manufacturer’s instructions. Fresh SkM01 medium was added and the plate was plate in the Incucyte® Live-Cell Analysis system for image acquisition in each well every hour for 24 hours.

### QUANTIFICATION AND STATISTICAL ANALYSIS

#### Immunofluorescence

Each hiPSC line was differentiated and imaged as 4 by 4 panels or individual pictures, and analyzed with the QuPath software. C-Met and Ecad-positive areas were determined by thresholding on the Alexa488 fluorescent channel (ECad: Threshold = 40 or regions manually drawn if low signal, C-Met: Threshold = 600, minimal object size= 2,500 µm²), and the number of nuclei was determined in the positive and negative areas using the Cell Detection tool with the following parameters: C-Met: DAPI threshold = 100, object size between 10 and 400 µm², background radius = 8 µm, sigma = 1.5 µm; E-Cad: DAPI threshold = 100, object size between 75 and 2000 µm², background radius = 8 µm, sigma = 2 µm. E-Cad pictures were acquired at higher magnification, explaining why different parameters had to be used. MF20- and a-actinin-positive areas were determined by thresholding on the TRITC channel (MF20: threshold = 100, α-actinin: threshold = 1000). Data were represented and analyzed with GraphPad PRISM 8.0.1. Non-parametric statistical tests were used to compare groups as sample size were low and we could not assume gaussian distributions nor homoscedasticity. The Mann-Whitney statistics was used when only two groups were compared, and the Kruskal-Wallis statistics for comparison of the 3 groups.

#### Cell migration assay

Data processing and determination of the relative wound density over time was performed with the dedicated Incucyte® analysis module. As previously mentioned, two independent experiments were performed. Each experiment included Day 7 progenitors derived from 6 hiPSC lines (3 Healthy and 3 DMD), each seeded in 8 wells of the Imagelock 96-well plate. Each hour, data points were collected in each well, and the RWD of each line was determined by averaging the data of the 8 wells.

## KEY RESOURCES TABLE

| REAGENT or RESOURCE | SOURCE | IDENTIFIER |
| --- | --- | --- |
| Antibodies |  |  |
| Myosin heavy chain | DSHB | Cat# MF20<br>RRID:AB_2147781 |
| alpha-Actinin (Sarcomeric) | Sigma-Aldrich | Cat# A7811;<br>RRID:AB_476766 |
| CD324 (E-cadherin) Alexa Fluor™ 488 | Thermo Fisher | Cat# 53-3249-80<br>RRID:AB_10671270 |
| Vimentin eFluor™ 570 | Thermo Fisher | Cat# 41-9897-80<br>RRID:AB_11220476 |
| Human HGFR/c-MET | R&D Systems | Cat# AF276<br>RRID:AB_355289 |
| Goat anti-Mouse IgG2b Alexa Fluor™ 555 | Thermo Fisher | Cat# A-21147<br>RRID:AB_2535783 |
| Donkey anti-Goat IgG Alexa Fluor™ 488 | Thermo Fisher | Cat# A-11055<br>RRID:AB_2534102 |
| Dystrophin | Leica | Cat# NCL-DYS2<br>RRID:AB_442081 |
| GAPDH | Novus | Cat# NB300-320<br>RRID:AB_10001796 |
| Alpha tubulin | Sigma-Aldrich | Cat# T5168<br>RRID:AB_477579 |
| Goat Anti-Mouse Immunoglobulins/HRP | Agilent | Cat# P0447<br>RRID:AB_2617137 |
| Polyclonal Rabbit Anti-Goat Immunoglobulins/HRP | Agilent | Cat# P0449<br>RRID:AB_2617143 |
| Chemicals, peptides, and recombinant proteins |  |  |
| mTeSR™ Plus | STEMCELL Technologies | Cat# 100-0276 |
| Matrigel® Matrix | Corning | Cat# 354234 |
| Versene solution | Thermo Fisher | Cat# 15040066 |
| CryoStor CS10 cryopreservation medium | STEMCELL Technologies | Cat# 07959 |
| DPBS | Thermo Fisher | Cat# 14190144 |
| Y-27632 ROCK inhibitor | Selleckchem | Cat# S1049 |
| DMEM/F12 | Thermo Fisher | Cat# 11320033 |
| CHIR99021 | STEMCELL Technologies | Cat# 72054 |
| StemMACS™ LDN-193189 | Miltenyi Biotec | Cat# 130-103-925 |
| Recombinant Human FGF2 | R&D Systems | Cat# 233-FB-010 |
| Recombinant Human IGF-1 | R&D Systems | Cat# 291-G1-200 |
| Recombinant Human HGF | R&D Systems | Cat# 294-HG-005 |
| KnockOut™ serum replacement | Thermo Fisher | Cat# 10828010 |
| Collagenase, type IV | Thermo Fisher | Cat# 1710419 |
| Trypsin-EDTA | Sigma-Aldrich | Cat# T4174 |
| Skeletal Muscle. Induction Medium | amsbio | Cat# SKM01 |
| Skeletal Muscle. Myoblast Medium | amsbio | Cat# SKM02 |
| Skeletal Muscle. Myotube Medium | Amsbio | Cat# SKM03 |
| Cultrex Rat Collagen I | R&D Systems | Cat# 3440-100-01 |
| Paraformaldehyde | Thermo Fisher | Cat# #28906 |
| Triton 100X | Eurobio | Cat# GAUTTR001 |
| BSA (immunofluorescence) | Sigma-Aldrich | Cat# A3059 |
| ProLong™ Gold Antifade reagent | Thermo Fisher | Cat# P36934 |
| Tris base | Sigma-Aldrich | Cat# 10708976001 |
| cOmplete™ Protease Inhibitor Cocktail | Merck | Cat# 11873580001 |
| Protease inhibitor cocktail | Sigma-Aldrich | Cat# P8340 |
| Igepal CA-630 | Sigma-Aldrich | Cat# I3021 |
| SDS solution | Sigma-Aldrich | Cat# 05030 |
| NuPAGE™ LDS sample buffer | Thermo Fisher | Cat# NP0007 |
| Dithiothreitol | Thermo Fisher | Cat# D1532 |
| NuPAGE™ Tris-acetate Gel | Thermo Fisher | Cat# EA03785BOX |
| NuPAGE™ Tris acetate SDS buffer | Thermo Fisher | Cat# LA0041 |
| Tween 20 | Merck | Cat# 8170721000 |
| Pierce™ ECL Western Blotting Substrate | Thermo Fisher | Cat# 32106 |
| Nuclease free water | Ambion | Cat# AM9932 |
| Methanol | Fisher Scientific | Cat# 10285131 |
| SUPERase In RNase inhibitor | Invitrogen | Cat# AM2694 |
| BSA (single-cell RNA-Seq) | New England Biolabs | Cat# B9000 |
| 1M Tris-HCl pH 7.5 | Thermo Fisher | Cat# 15567027 |
| 5M NaCl (single-cell RNA-Seq) | Thermo Fisher | Cat# AM9759 |
| 1M MgCl<sub>2</sub> | Thermo Fisher | Cat# AM9530G |
| IGEPAL CA-360 | Sigma-Aldrich | Cat# 18896 |
| Diethyl pyrocarbonate | Sigma-Aldrich | Cat# D5758 |
| 10mM dNTP | Thermo Fisher | Cat# 18427013 |
| Superscript IV reverse transcriptase | Thermo Fisher | Cat# 18090200 |
| RNaseOUT Recombinant Ribonuclease Inhibitor | Thermo Fisher | Cat#10777019 |
| 0.5M EDTA pH 8.0 | Thermo Fisher | Cat# 15575020 |
| Spermidine | MP Biomedicals | Cat# 0219485201 |
| Elution Buffer | Qiagen | Cat# 19086 |
| DAPI | Thermo Fisher | Cat# D1306 |
| DNA binding buffer | Zymo Research | Cat# D4003-1 |
| AMPure XP beads | Beckman Coulter | Cat# A63880 |
| Ethanol | Decon Labs | Cat# 2716 |
| NEBNext® High Fidelity 2X PCR Master Mix | New England Biolabs | Cat# M0541 |
| Novex® TBE-PAGE gels, 6% | Thermo Fisher | Cat# EC6265BOX |
| Quick-Load 2-log DNA Ladder | New England Biolabs | Cat# N0550 |
| Critical commercial assays |  |  |
| Nextseq 500/550 V2 75 cycle kit | illumina | Cat# 20024906 |
| DC Protein Assay | Bio-Rad | Cat# 5000111 |
| Tagment DNA TDE1 Enzyme and Buffer Kit | illumina | Cat# 20034197 |
| Evercode™ Whole Transcriptome Mini Kit | Parse Biosciences | Cat# EC-W01010 |
| Cell Fixation Kit | Parse Biosciences | Cat# SB1001 |
| Qubit dsDNA HS Kit | Thermo Fisher | Cat# Q32851 |
| NEBNext® Ultra™ II Non-Directional RNA Second Strand Synthesis Module | New England Biolabs | Cat# E6111 |
| Deposited data |  |  |
| sciRNA-Seq myogenesis time course data | This paper | GEO accession number: GSE233605 |
| Bulk RNA Seq Day 10 data | Mournetas *et al.*<sup>1</sup> | ArrayExpress: E-MTAB-8321 |
| Split-pool barcoding scRNASeq Day 10 data | This paper | GEO accession number: GSE233606 |
| Supplementary Tables S1 to S7 | This paper | Zenodo DOI: 10.5281/zenodo.10849178 |
| Full Western blots films related to Figure S2A | This paper | Zenodo DOI: 10.5281/zenodo.7937310 |
| Immunofluorescence pictures related to Figure 3A-B | This paper | Zenodo DOI: 10.5281/zenodo.8325281 |
| Immunofluorescence pictures related to Figure 4B-C | This paper | Zenodo DOI: 10.5281/zenodo.7950222 |
| Immunofluorescence pictures related to Figure 4D-E | This paper | Zenodo DOI: 10.5281/zenodo.8325290 |
| Immunofluorescence pictures related to Figure 5A-B | This paper | Zenodo DOI: 10.5281/zenodo.8325317 |
| Experimental models: Cell lines |  |  |
| UC3-4 Healthy hiPSC line, passage 32 | Laboratory of David L Mack | N/A |
| DMD 72039 D3 hiPSC line, passage 22 | Laboratory of David L Mack | N/A |
| UC3-4 DMD CRISPR hiPSC line, passage 41 | Laboratory of David L Mack | N/A |
| NIH/3T3 cell spike (single-cell RNA-Seq) | ATCC | N/A |
| Oligonucleotides |  |  |
| sciRNA-Seq barcoded oligonucleotides (see Table S1) | IDT | N/A |
| Custom P5 and P7 primers (see Table S1) | IDT | N/A |
| Software and algorithms |  |  |
| Fiji (imageJ) | Schindelin *et al.*<sup>2</sup> | https://fiji.sc/ |
| QuPath version 0.4.3 | Bankhead *et al.*<sup>3</sup> | https://qupath.github.io/ |
| PRISM version 8 | Graphpad | https://www.graphpad.com/features |
| bcl2fastq version 2.16.0.10 | illumina | https://emea.support.illumina.com/sequencing/sequencing_software/bcl2fastq-conversion-software.html |
| trim_galore version 0.4.5 | Felix Krueger | https://www.bioinformatics.babraham.ac.uk/projects/trim_galore/ |
| STAR version 2.5.2b | Dobin *et al.*<sup>4</sup> | https://github.com/alexdobin/STAR |
| R version 4.2.2 | https://cran.r-project.org/ |  |
| dplyr version 1.1.0 | Wickham *et al.* | https://dplyr.tidyverse.org/ |
| monocle version 1.3.1 | Cao *et al.*<sup>5,6</sup> | https://cole-trapnell-lab.github.io/monocle3/ |
| ggplot2 version 3.1.4 | Wickham *et al.* | https://ggplot2.tidyverse.org/ |
| sci-RNA-Seq and split-pool barcoding analysis pipelines | This paper | Zenodo<br>DOI :<br>10.5281/zenodo.8348000 |

---

**Supplemental Table S1:** cell_data file obtained after the initial sciRNA-Seq experiment. The genotype, time point, number of expressed genes and cluster of origin are indicated for each individual cell.

**Supplemental Table S2:** Differentially expressed genes (p-adj < 0.0001) obtained after branch expression analysis modeling (BEAM) and the cluster to which they belong on the heatmap shown in Figure 1F.

**Supplemental Table S3:** Differentially expressed genes (p-adj < 0.01) detected at Day 10 after sciRNA-Seq.

**Supplemental Table S4:** Details of the differentiation experiments performed to obtain Figures 1 to 5.

**Supplemental Table S5:** Differentially expressed genes (p-adj < 0.01) detected at Day 10 after split-pool barcoding RNA-Seq.

**Supplemental Table S6:** Differentially expressed genes (abs(log2(fold-change)) > 1 & p-adj < 0.01) detected at Day 10 after bulk RNA-Seq in the study previously published by our group^20^.

**Supplemental Table S7:** List of barcoded primers used in the RT reaction and in the illumine library preparation of the sciRNA-Seq experiment.

The Supplementary Tables S1 to S7 have been deposited on the Zenodo database, and are available using the following DOI: 10.5281/zenodo.10849178

**Supplemental Figure S1:**
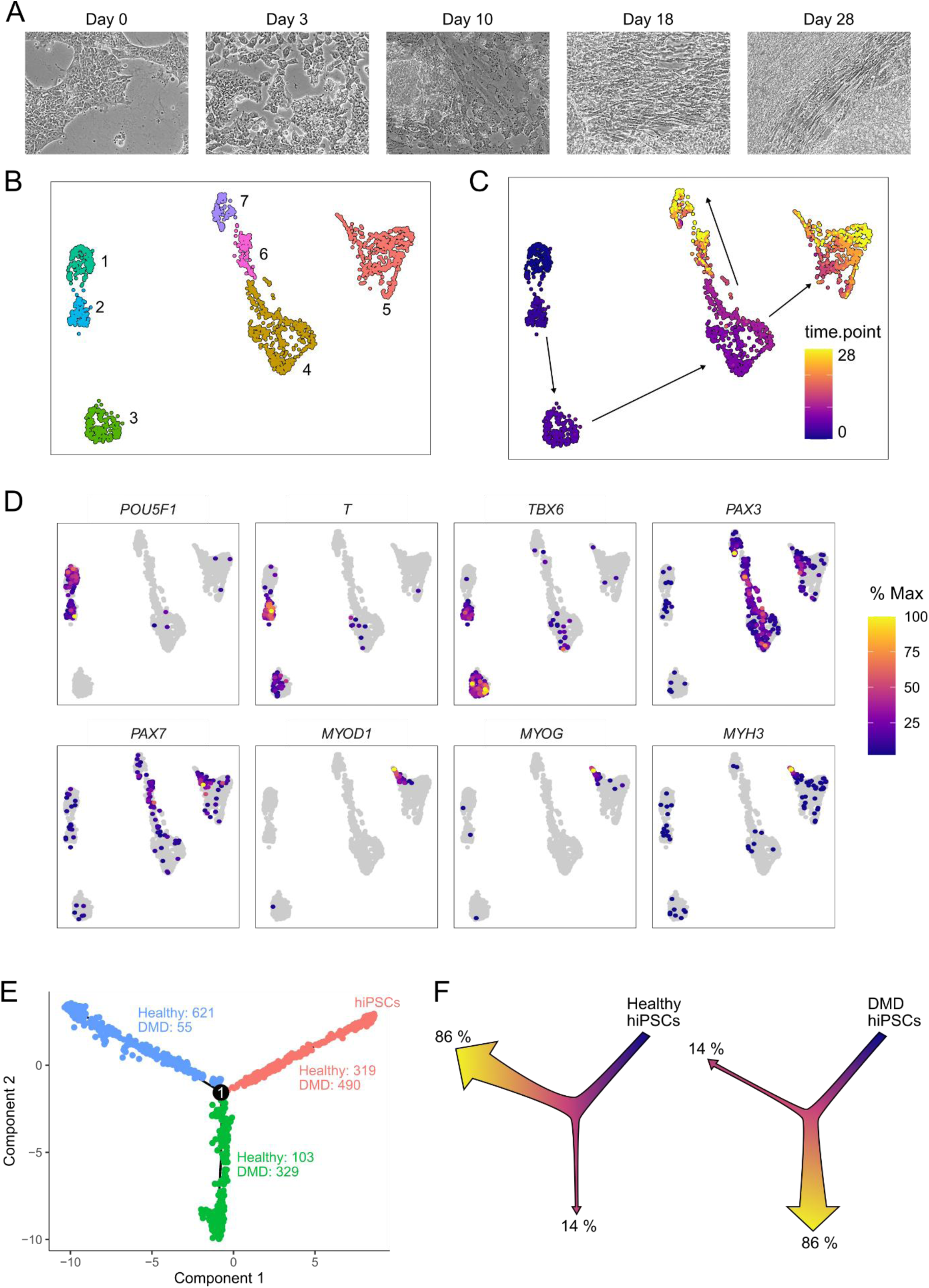
Myogenic differentiation of hiPSCs at the single-cell resolution. (A) Observation of hiPSC cultures along the differentiation at process by phase contrast microscopy. (B) UMAP plot showing the 1917 individual cells colored by cluster. (C) UMAP plot showing the 1917 individual cells colored by collection time point from Day 0 to D28. (D) Expression of successive developmental markers in hiPSCs along the myogenic differentiation. (E) Number of DMD and Healthy cells on the three branches of the developmental trajectory. (F) Schematic representation of the developmental trajectory followed by healthy (left panel) and DMD hiPSCs (right panel).

**Supplemental Figure S2:**
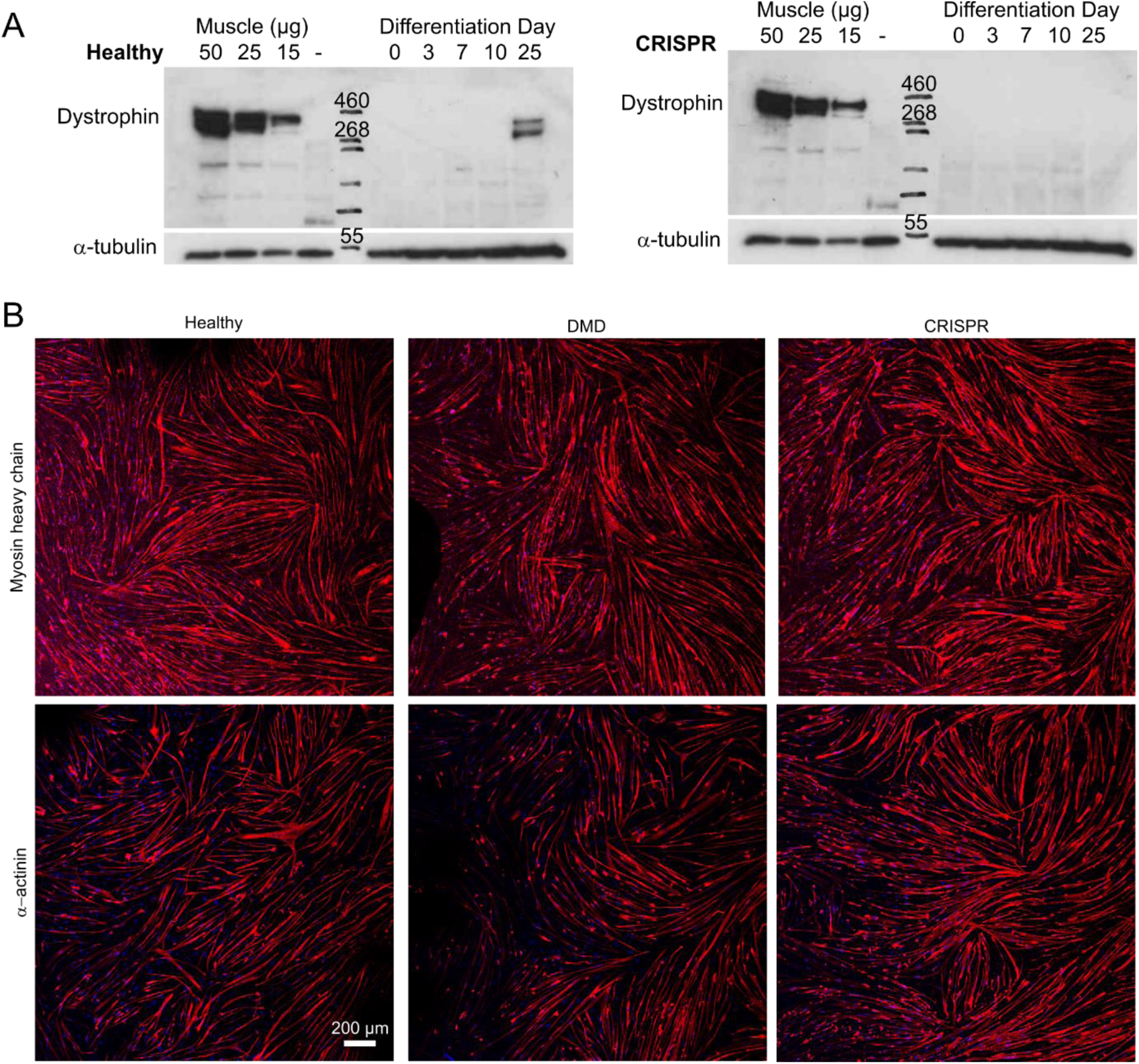
(A) Expression of dystrophin along the myogenic differentiation of CRISPR hiPSCs (right panel) and healthy control cells (left panel). Proteins were extracted from cell samples collected from Day 0 to Day 25 or from the pectoral muscles of a healthy rat as positive controls (“Muscle”) and a Dmd*^mdx^* rat as a negative control (“-”), and subjected to anti-dystrophin western-blot analysis. Alpha tubulin (a-tubulin) was used as a control. (B) Fluorescent staining of myosin heavy chains and a-actinin (red) in myotubes derived from Healthy, DMD and CRISPR hiPSCs. Scale bar = 200 µm.

**Supplemental Figure S3:**
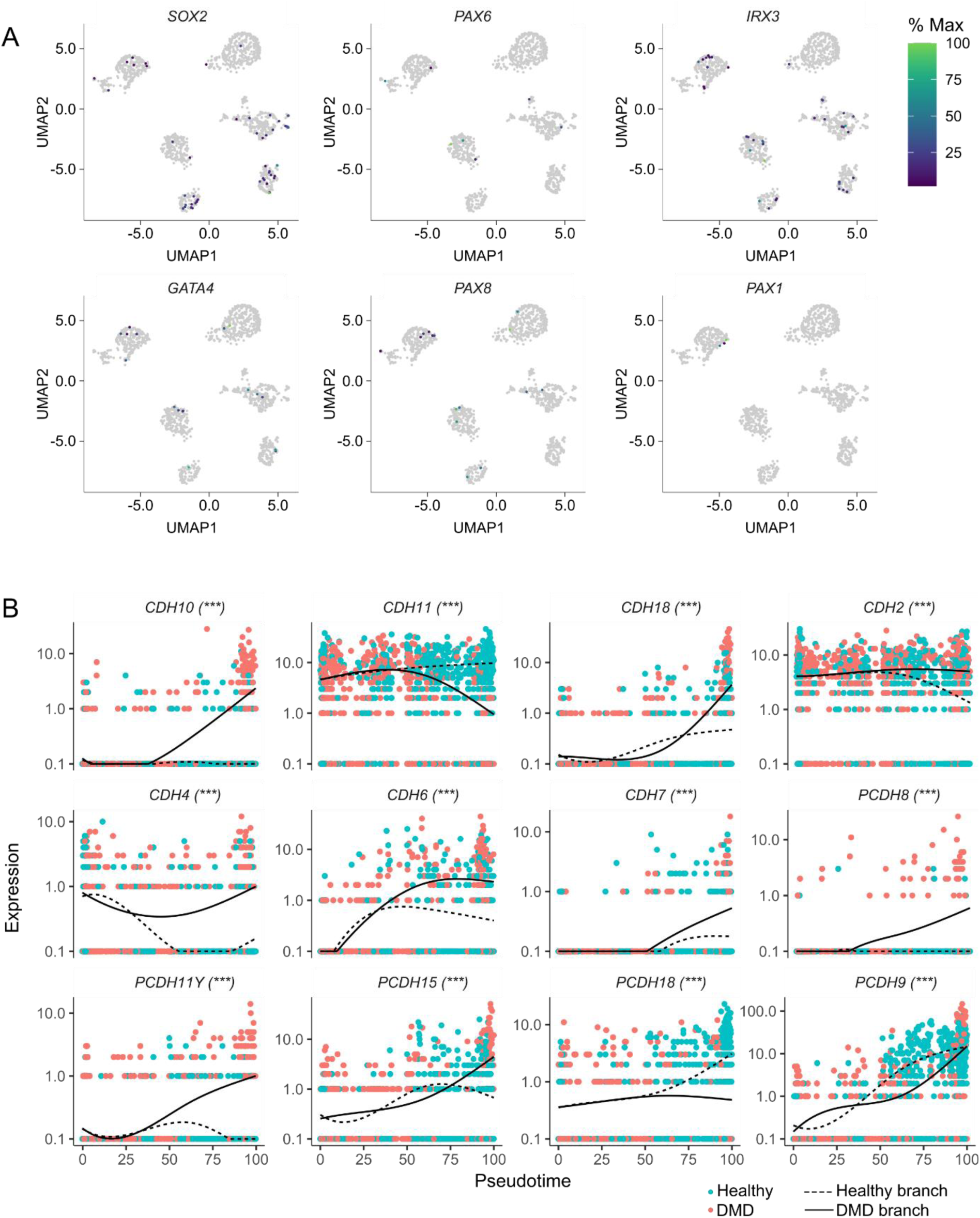
(A) Expression of developmental markers from alternative lineages at Day 10 (neural tube: *SOX2*, *PAX6*, *IRX3*, lateral plate mesoderm: *GATA4*, intermediate mesoderm: *PAX8* and sclerotome: *PAX1*). (B) Pseudotime expression of *CDH* and *PCDH* along the Healthy and the DMD branches in the sci-RNA-Seq data set. Individual cells are colored by their hiPSC line of origin and ordered along the pseudotime axis. BEAM statistics: ***: adjusted p-value < 0.001.

**Supplemental Figure S4:**
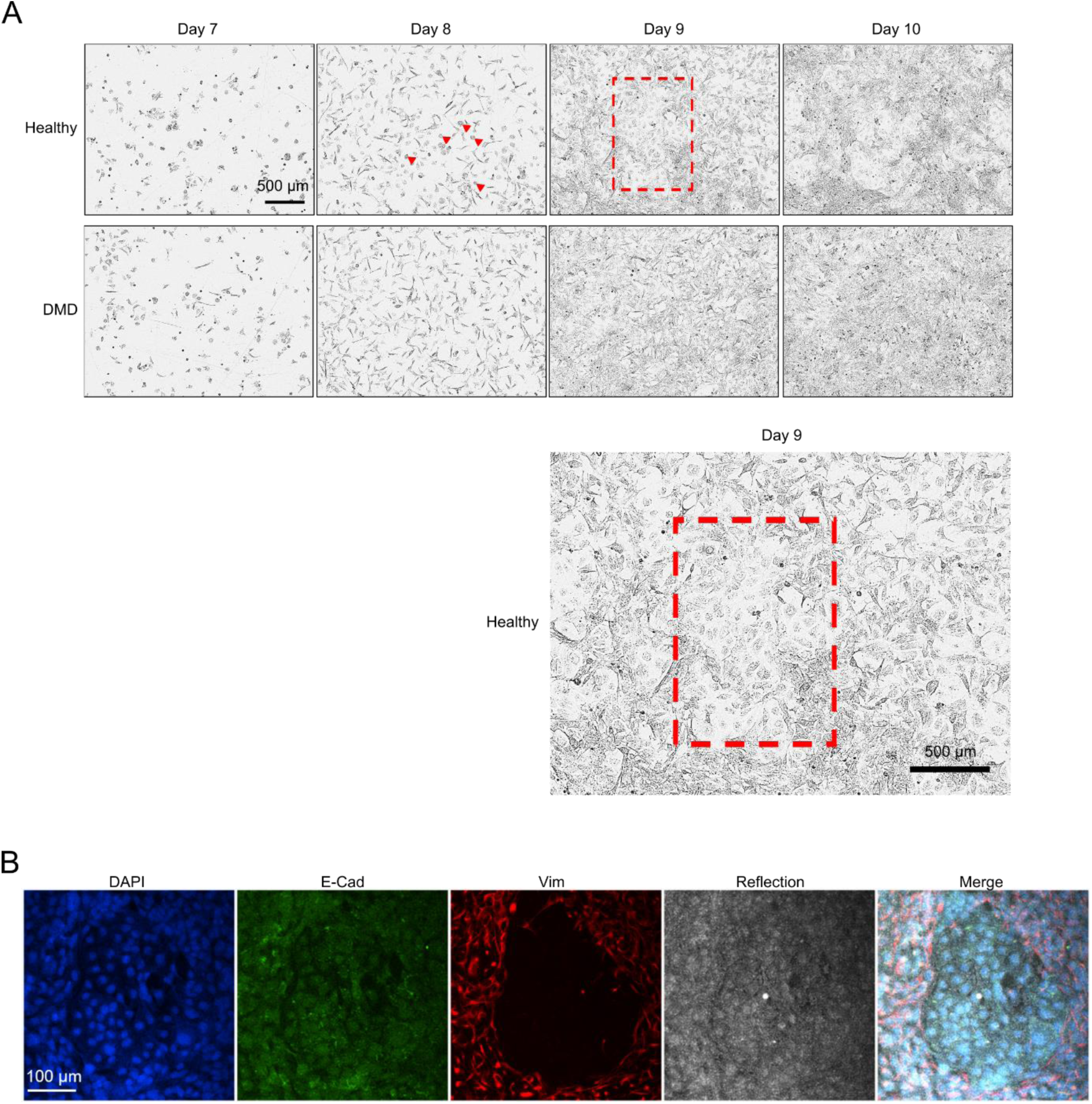
(A) Evolution of hiPSC cultures between Day 7 and Day 10 of the myogenic differentiation protocol. Red arrowheads indicate “round” cells appearing from Day 8 and at the core of the epithelial islets visible from Day 9 – Day 10 (red rectangle). (B) Details of the immunofluorescence staining with the addition of reflection imaging on the confocal microscope, which helps specify that the flattened cells correspond to the Vim-negative epithelial islets.

**Supplemental Figure S5:**
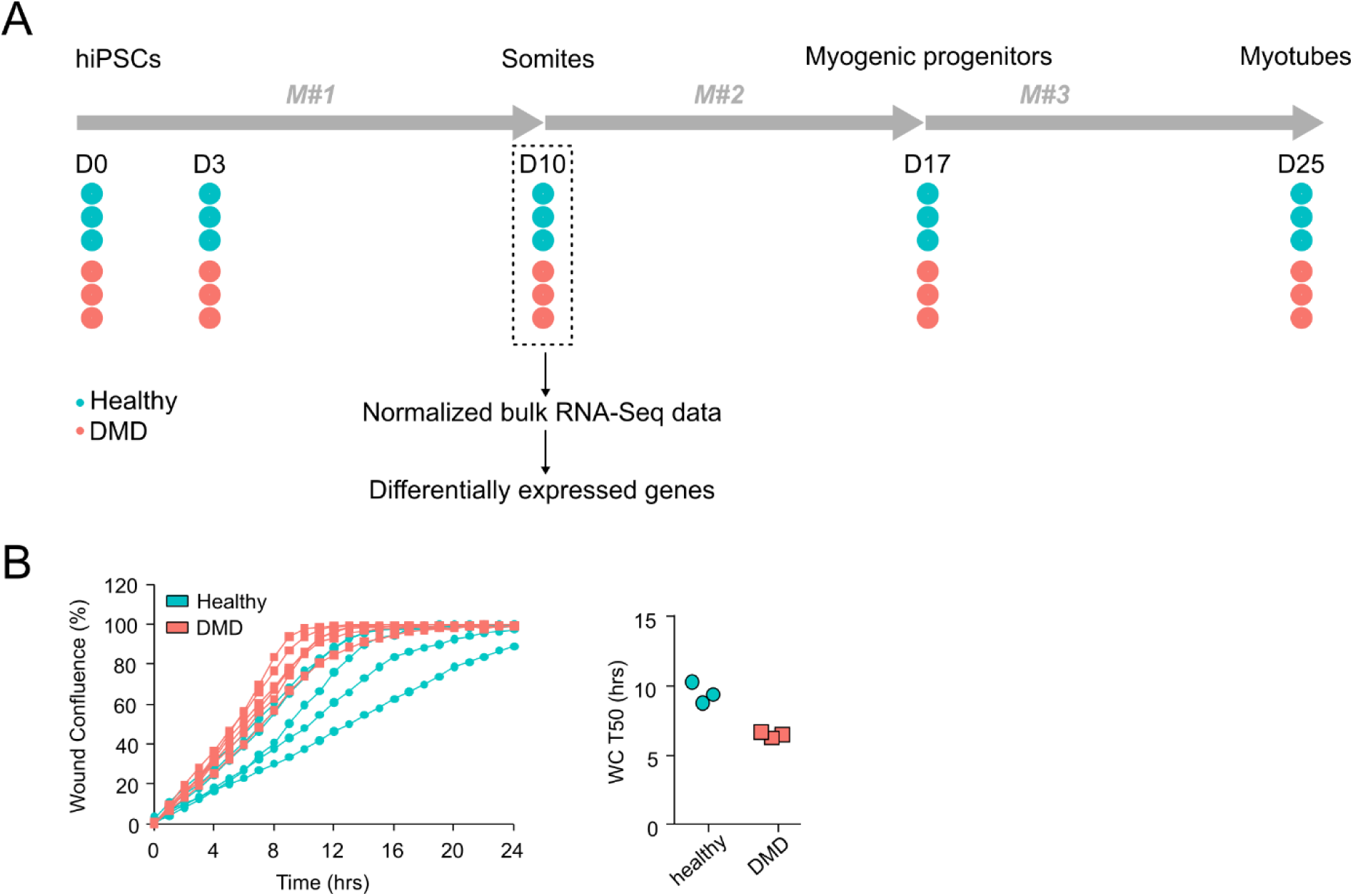
(A) Differentiation of DMD and Healthy hiPSCs and cell collection protocol in the analyzed study^20^. Each individual line is indicated by a blue (Healthy) or a red (DMD) dot, and the analysis at Day 10 is highlighted by a dotted square. (B) Monitoring of the wound confluence (WC) over time in the 3 DMD and 3 Healthy control lines. For each line, the time to reach a confluence of 50% in the wound area (WC T50) was determined and averaged from 2 x 8 wells per line in 2 independent experiments.

